# Bro1 stimulates Vps4 activity to promote Intralumenal Vesicle Formation during Multivesicular Body biogenesis

**DOI:** 10.1101/2020.07.27.223255

**Authors:** Chun-Che Tseng, Shirley Dean, Brian A. Davies, Ishara F. Azmi, Natalya Pashkova, Johanna A. Payne, Jennifer Staffenhagen, Matt West, Robert C. Piper, Greg Odorizzi, David J. Katzmann

**Author notes:** These authors contributed equally to these studies (Chun-Che Tseng, Shirley Dean, Brian A. Davies).

## Abstract

Endosomal sorting complexes required for transport (ESCRT-0, -I, -II, -III) execute cargo sorting and intralumenal vesicle (ILV) formation during conversion of endosomes to multivesicular bodies (MVBs). The AAA-ATPase Vps4 regulates the ESCRT-III polymer to facilitate membrane remodeling and ILV scission during MVB biogenesis. Here we show that the conserved V domain of ESCRT-associated protein Bro1 (the yeast homolog of mammalian proteins ALIX and HD-PTP) directly stimulates Vps4. This activity is required for MVB cargo sorting. Furthermore, the Bro1 V domain alone supports Vps4/ESCRT-driven ILV formation *in vivo* without efficient MVB cargo sorting. These results reveal a novel activity of the V domains of Bro1 homologs in licensing ESCRT-III-dependent ILV formation and suggest a role in coordinating cargo sorting with membrane remodeling during MVB sorting. Moreover, ubiquitin binding enhances V domain stimulation of Vps4 to promote ILV formation via the Bro1/Vps4/ESCRT-III axis, uncovering a novel role for ubiquitin during MVB biogenesis in addition to facilitating cargo recognition.

**Summary:** Cargo sorting is coordinated with intralumenal vesicle budding during ESCRT-mediated multivesicular body biogenesis. Bro1 V domain stimulates Vps4 to promote ESCRT-III-driven intralumenal vesicle formation in a manner required for this coordinated process.

## Introduction

The Endosomal Sorting Complexes Required for Transport (ESCRTs) have been implicated in a number of cellular membrane remodeling processes: intralumenal vesicle (ILV) generation during the conversion of endosomes into multivesicular bodies (MVBs, relevant to both lysosomal degradation and exosome biogenesis; Baietti et al., 2012, reviewed by Hanson and Cashikar, 2012), abscission during cytokinesis (reviewed by Vietri et al., 2020), viral budding (Votteler and Sundquist, 2013), nuclear pore surveillance (Webster et al., 2014), autophagy (Lee et al., 2007a; Takahashi et al., 2018), and membrane repair (Skowyra et al., 2018). These studies have highlighted the conserved role of ESCRT-III and associated factors in membrane deformation. ESCRT-III-dependent membrane remodeling is typically coordinated with upstream events, such as cargo recognition in MVB sorting (reviewed by Piper and Lehner, 2011; Williams and Urbe, 2007). Ubiquitylated cargoes destined for lysosomal destruction are actively recognized by a host of ubiquitin (Ub)-binding domains within ESCRTs -0, -I, and -II as well as in the Bro1 Domain Family Proteins, and sequestration of these ubiquitylated cargoes form microdomains at which ESCRT-II activates ESCRT-III polymerization to drive ILV formation (reviewed by Schmidt and Teis, 2012; Williams and Urbe, 2007). In yeast depletion of Ub from the site of MVB sorting precludes cargo sorting as well as ILV formation itself, indicating a level of coordination between cargo recognition and ESCRT-III-mediated ILV generation (MacDonald et al., 2012a; Stringer and Piper, 2011). ESCRT-III assembly, remodeling and disassembly drive membrane budding, although how this cycle is regulated to coordinate cargo transfer into ILVs remains an unresolved question.

ESCRT-III subunits are monomeric in the cytosol and undergo an ordered polymerization on membranes into filamentous spirals responsible for membrane remodeling (reviewed by Schmidt and Teis, 2012). ESCRT-III function is intimately connected to the AAA-ATPase Vps4, which supports both 1) dynamic exchange of subunits during ESCRT-III polymerization (Adell et al., 2017; Mierzwa et al., 2017; Pfitzner et al., 2020) and 2) ESCRT-III disassembly at the completion of the reaction (Adell and Teis, 2011; Babst et al., 1997; Babst et al., 1998; Davies et al., 2010). A series of regulators including ESCRT-III itself, the Vps4 co-factor Vta1/LIP5, and Bro1 serve to coordinate the activities of ESCRT-III and Vps4 to optimize ESCRT function during MVB sorting (Azmi et al., 2006; Azmi et al., 2008; Merrill and Hanson, 2010; Shim et al., 2008; Wemmer et al., 2011). How these factors act in concert during MVB sorting is unclear. The Bro1 Domain Family members, including mammalian ALIX/PDCD6IP and HD-PTP/PTPN23 and the aforementioned yeast Bro1, make multiple contributions to ESCRT-mediated events. This family is characterized by the presence of: the amino-terminal Bro1 Domain (BOD) that binds the ESCRT-III subunit CHMP4/Snf7 (Lee et al., 2016; McCullough et al., 2008; Wemmer et al., 2011); the middle V domain - two helix bundles that form two arms of a V-shaped structure capable of interacting with both Ub and YPX_n_L motifs found within Gag, Syntenin, and Rfu1 (Baietti et al., 2012; Carlton et al., 2008; Kimura et al., 2014); and the carboxyl-terminal Proline Rich Region (PRR) that both facilitates associations with ESCRT-I and other factors (e.g. Ub isopeptidase Doa4 interaction with yeast Bro1 PRR) and, in the context of ALIX, auto-inhibits the BOD and V domain (Buysse et al., 2020; Johnson et al., 2017; Luhtala and Odorizzi, 2004; Nikko and André, 2007; Richter et al., 2013; Zhai et al., 2011). Bro1 Domain Family members contribute to 1) Ub-dependent and -independent cargo recognition in concert with or in parallel to the early ESCRTs and 2) regulating ESCRT-III dynamics by facilitating CHMP4/Snf7 activation and inhibiting Vps4 disassembly of ESCRT-III. These diverse contributions may suggest Bro1 Domain Family members serve roles coordinating cargo entry into budding ILVs.

Loss of Bro1 severely diminishes ESCRT-dependent functions and disrupts ILV formation as does loss of the ESCRT-III subunit Snf7 (Babst et al., 2002; Odorizzi et al., 2003). However, when the interaction of Snf7 with Bro1 BOD is disrupted, ILV formation is still observed albeit at a reduced level (Wemmer et al., 2011), suggesting that other features of and interactions with Bro1 drive ILV formation. Here we find that Bro1 V domain supports ESCRT-dependent MVB biogenesis *in vivo* through a process that uncouples cargo sorting from vesicle formation. The V domain interacts with Vps4 MIT domain and stimulates Vps4 ATPase activity *in vitro* in a manner regulated by Bro1V binding Ub. Stimulation of Vps4 is critical for Bro1 function since V-domain mutations that preserve Vps4 and Ub-binding but specifically perturb the ability to stimulate Vps4 ATPase activity showed defects in MVB cargo sorting and ILV formation *in vivo*. These results indicate that ESCRT-driven ILV formation is separable from cargo sorting and suggest that Bro1 Domain Family members coordinate ubiquitylated cargo sorting into ILVs whilst promoting ILV formation via the Vps4/ESCRT-III axis.

## Results

### Bro1 V domain promotes ESCRT-dependent ILV formation

The BOD of Bro1 interacts with the ESCRT-III subunit Snf7 (Kim et al., 2005) to promote ESCRT-III polymerization through both facilitating Snf7 activation (Tang et al., 2016) and modulating Vps4-mediated disassembly of ESCRT-III (Wemmer et al., 2011). Mutation of Bro1 or Snf7 disrupting their association abrogates MVB cargo sorting (Kim et al., 2005; Wemmer et al., 2011), however a Snf7 mutant defective for BOD binding [Snf7(L231A, L234A)] retained the capacity to support ILV formation (Wemmer et al., 2011), implying other domains of Bro1 are critical to drive ILV formation. To explore this idea, we examined the phenotype of expressing Bro1 lacking its N-terminal BOD (*bro1*^Δ*BOD*^ aa370-844) as the sole allele of *BRO1*. The *bro1*Δ cells expressing *bro1*^Δ*BOD*^ under the control of *TEF1* promoter displayed spherical endosomal structures containing ILVs (Figure 1A, Movie S1). In contrast, cells lacking Bro1 altogether, *brol*Δ. were deficient in ILVs and displayed multi-lamellar endosomal stacks characteristic of “class E vps” endosomal compartments ((Luhtala and Odorizzi, 2004) and Figure 1A, Movie S2; a rare *bro1*Δ cell with ILVs is shown in Figure S1A). Compared to WT cells *bro1*^Δ*BOD*^ cells showed reductions in ILVs per MVB, budding profile frequency and budding profile surface area (Figure 1B, 1C, Movie S3), however the sizes of ILVs were equivalent to those in WT cells (Figure 1C). While aspects of the *bro1*^Δ*BOD*^ MVBs differed from those in WT cells, the striking result was their presence in *bro1*^Δ*BOD*^ cells in contrast to their absence in *bro1*Δ cells (Figure 1B). This result indicates that Bro1 BOD is not required to support ILV formation, consistent with the Snf7(L231A, L234A) analysis (Wemmer et al., 2011).

**FIGURE 1.**
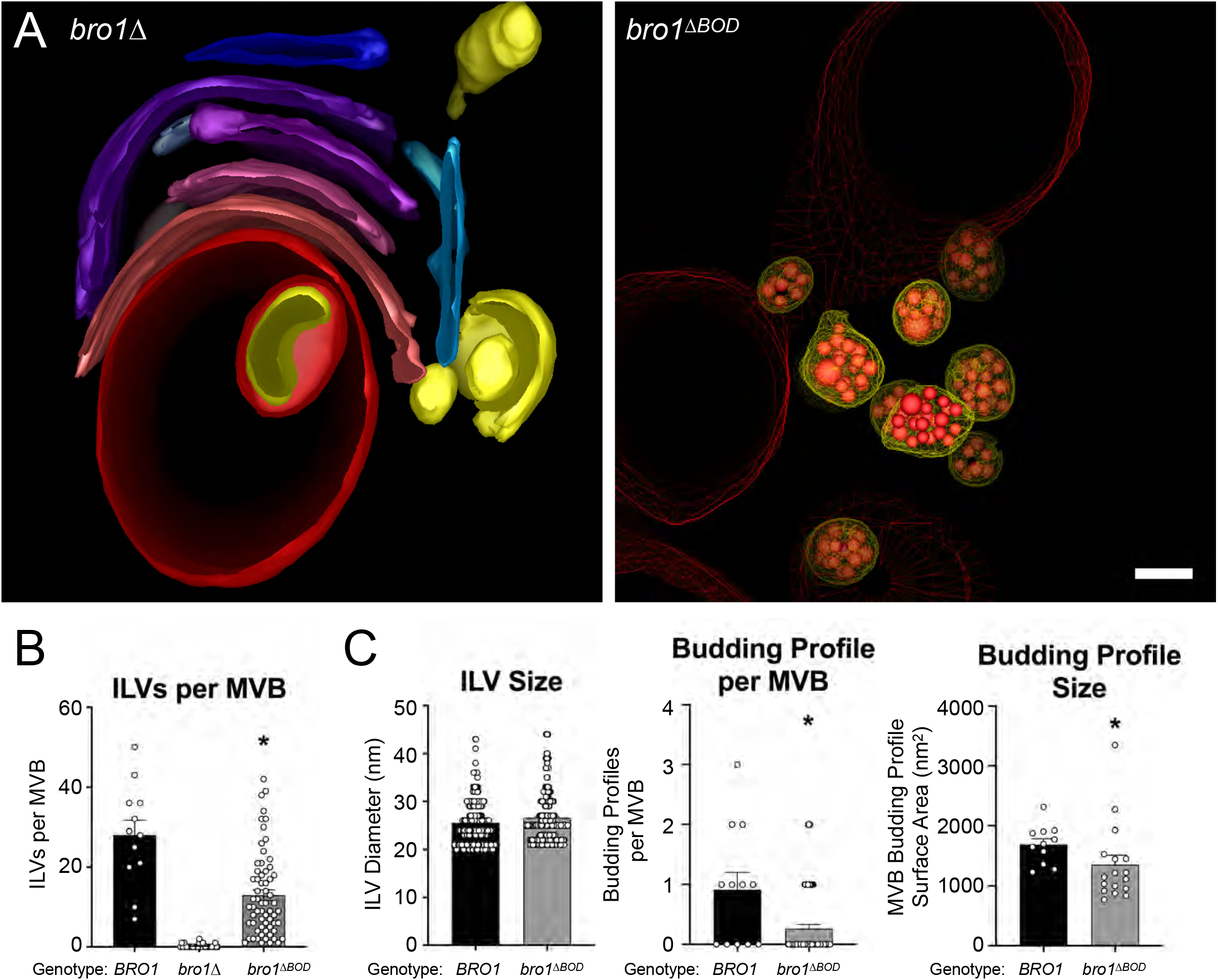
Bro1^ΔBOD^ supports ILV formation. Three-dimensional models reconstructed from 200-nm-thick section electron tomograms of *bro1*Δ (GOY65) and *bro1*^Δ*BOD*^ (*bro1*Δ::*TEF1*p-*bro1*Δ*BOD;* CTY2) cells. The *bro1*Δ cells have class E compartments, which are flattened stacks of endosomal membranes that generally lack internal vesicles; these stacks are shown in different colors to differentiate individual membranes. For *bro1*Δ*BOD*, the limiting membrane of MVBs are labeled yellow, the ILVs are highlighted in red, and the vacuole limiting membrane is labeled as red mesh. Scale bar = 100nm. **(B)** WT (SEY6210), *bro1*Δ (GOY65), and *bro1*^Δ*BOD*^ (CTY2) were analyzed by electron tomography and quantified for number of ILV per MVB. * indicates statistically significant differences compared to WT and *bro1*Δ. (**C)** WT (SEY6210), and *bro1*Δ*BOD* (CTY2) were analyzed by electron tomography and quantified to assess ILV size (diameter), individual budding profile size (surface area) and the frequency of incomplete ILV budding events (budding profiles per MVB). A minimum of 13 MVBs and 337 ILVs from at least 10 cells were quantified. Error bars indicate S.E.M. * indicates statistically significant differences compared to WT.

Bro1 requires BOD interaction with Snf7 for MVB cargo sorting (Kim et al., 2005; Wemmer et al., 2011); however, we observed ILV formation upon expression of bro1^ΔBOD^. To determine whether Bro1^ΔBOD^ could mediate specific MVB cargo sorting, we examined the localization of MVB cargoes delivered from endocytic (Ste2, Mup1) and biosynthetic pathways (Cps1, Ub-Cps1, Cos5, Sna3) as well as a vacuolar limiting membrane protein (DPAP-B/Dap2) typically excluded from MVB sorting. Though *bro1*^Δ*BOD*^ cells clearly made MVBs, MVB cargo proteins failed to sort into MVBs and instead were localized to the limiting membrane of the vacuole and a perivacuolar endosomal compartments (Figure 2A). Immunoblot analysis of two MVB cargos, Sna3 and Mup1, with anti-GFP antibodies showed that a modest level of delivery to the endosomal lumen did occur as gauged by the presence of a ~24 kDa fragment of GFP that is produced in the vacuolar lumen in a Snf7-, Vps4-and Bro1-dependent manner (Figure 2B and C). These sorting defects in *bro1*^Δ*BOD*^ cells did not appear to be related to perturbed cargo ubiquitylation since immunoblotting of Sna3 and Mup1 showed that a substantial proportion of these cargos were ubiquitylated. To confirm that *bro1*^Δ*BOD*^ cells made ILVs, we also followed the sorting of NBD-Phosphatidylcholine (NBD-PC), a fluorescent lipid that partitions into ILVs in an ESCRT-dependent manner (Bilodeau et al., 2002; Hanson et al., 2002; Shields et al., 2009), and Figure 3A-C, Figure S1). Expression of Bro1^ΔBOD^ in *bro1*Δ cells resulted in a significant percentage of cells with lumenal NBD-PC sorting (77%), consistent with the ability of Bro1^ΔBOD^ to support ILV formation. Over-expression of full-length Bro1 under the control of the *TEF1* promoter (Figure S2A) reduced the percentage of cells with NBD-PC sorted into the vacuolar lumen (98% vs 85%; Figure 3C), consistent with the dominant negative effect of over-expressing Bro1 (Wemmer et al., 2011). Bro1^ΔBOD^-mediated NBD-PC sorting was dependent on Vps4 as well as ESCRT-0, ESCRT-I, ESCRT-II and ESCRT-III (Figure 3C, Figure S2B), indicating that Bro1^ΔBOD^ uses the expected ESCRT-dependent pathway for ILV formation. Moreover, expression of Bro1^ΔBOD^ under the control of *BRO1* promoter was able to support NBD-PC sorting (Figure S2E, S2F) albeit to a lesser degree than highly-expressed Bro1^ΔBOD^. Together, these results indicate that while Bro1^ΔBOD^ can support MVB biogenesis (e.g., *bro1*^Δ*BOD*^ cells exhibit ~50% ILVs per MVB compared to WT cells), Bro1^ΔBOD^ is unable to support efficient sorting of MVB cargos into ILVs.

**FIGURE 2.**
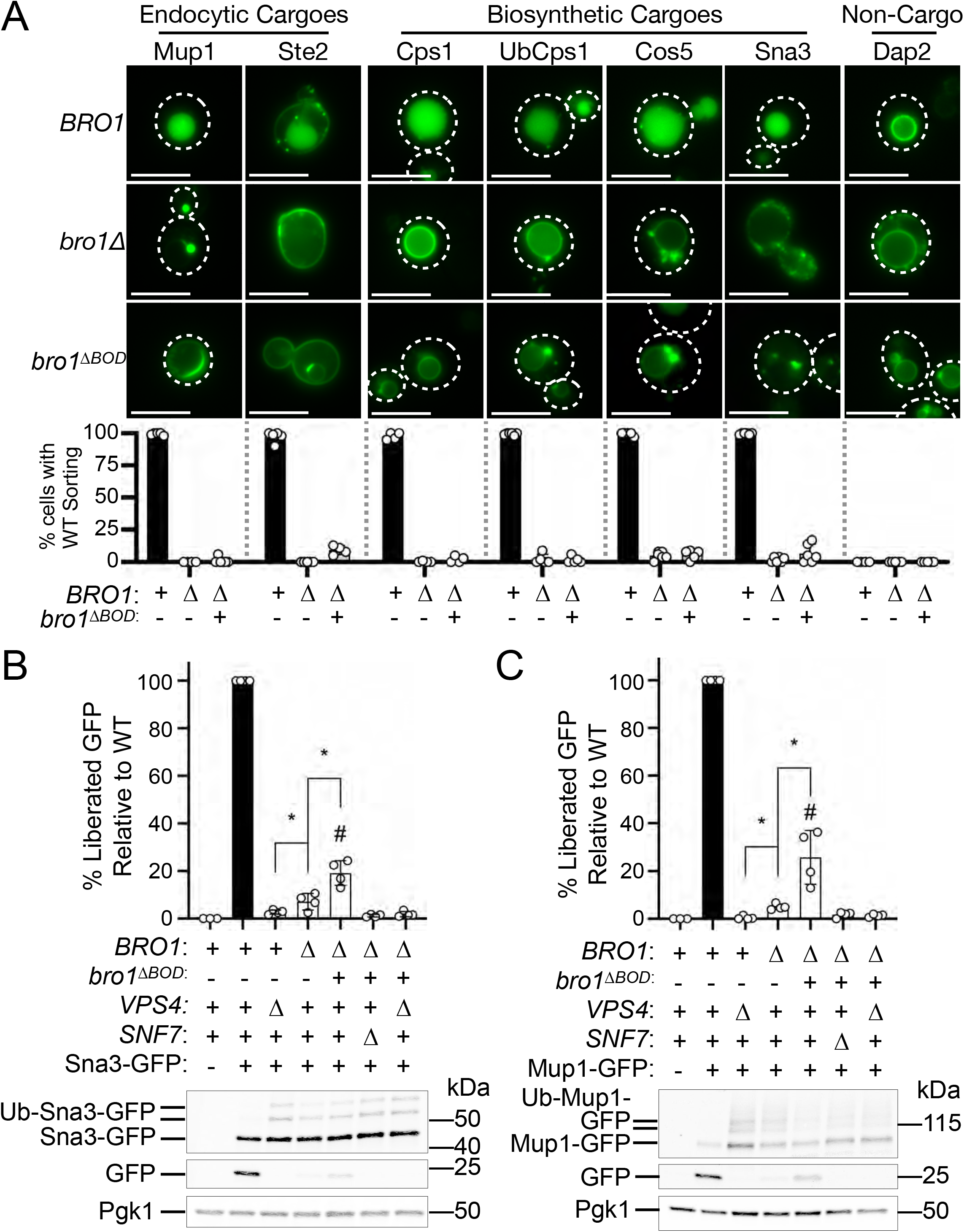
Bro1ΔBOD does not support efficient MVB cargo sorting. **(A)** WT (SEY6210), *bro1*Δ (GOY65), or *bro1*^Δ*BOD*^ (*bro1*Δ::*TEF1*p-*bro1*Δ*BOD*; CTY2) cells were transformed with the indicated GFP-tagged cargo plasmid to assess MVB sorting using live cell fluorescence microscopy. Percentage of cells with WT sorting signal was quantified and calculated from at least 183 cells from four independent experiments performed on four different days from three different transformations. White dashed lines indicate cell boundaries. Scale bar = 5μm. **(B-C)** Representative immunoblots showing expression levels of GFP-tagged cargo, liberated GFP and Pgk1 (loading control) using lysates of WT (SEY6210), *vps4*Δ (MBY3), *bro1*Δ (GOY65), *bro1*^Δ*BOD*^ (*bro1*Δ::*TEFL1*p-*bro1*^Δ*BOD*^; CTY2), *bro1*^Δ*BOD*^ *snf7*Δ (CTY12), or *bro1*^Δ*BOD*^ *vps4*Δ (CTY5) cells transformed with Sna3-GFP (B) or Mup1-GFP (C). Data was quantified from four independent experiments performed on four separate days from two transformations. Error bars indicate S.D. * indicates statistically significant difference. # indicates statistically significant difference compared to WT.

**FIGURE 3.**
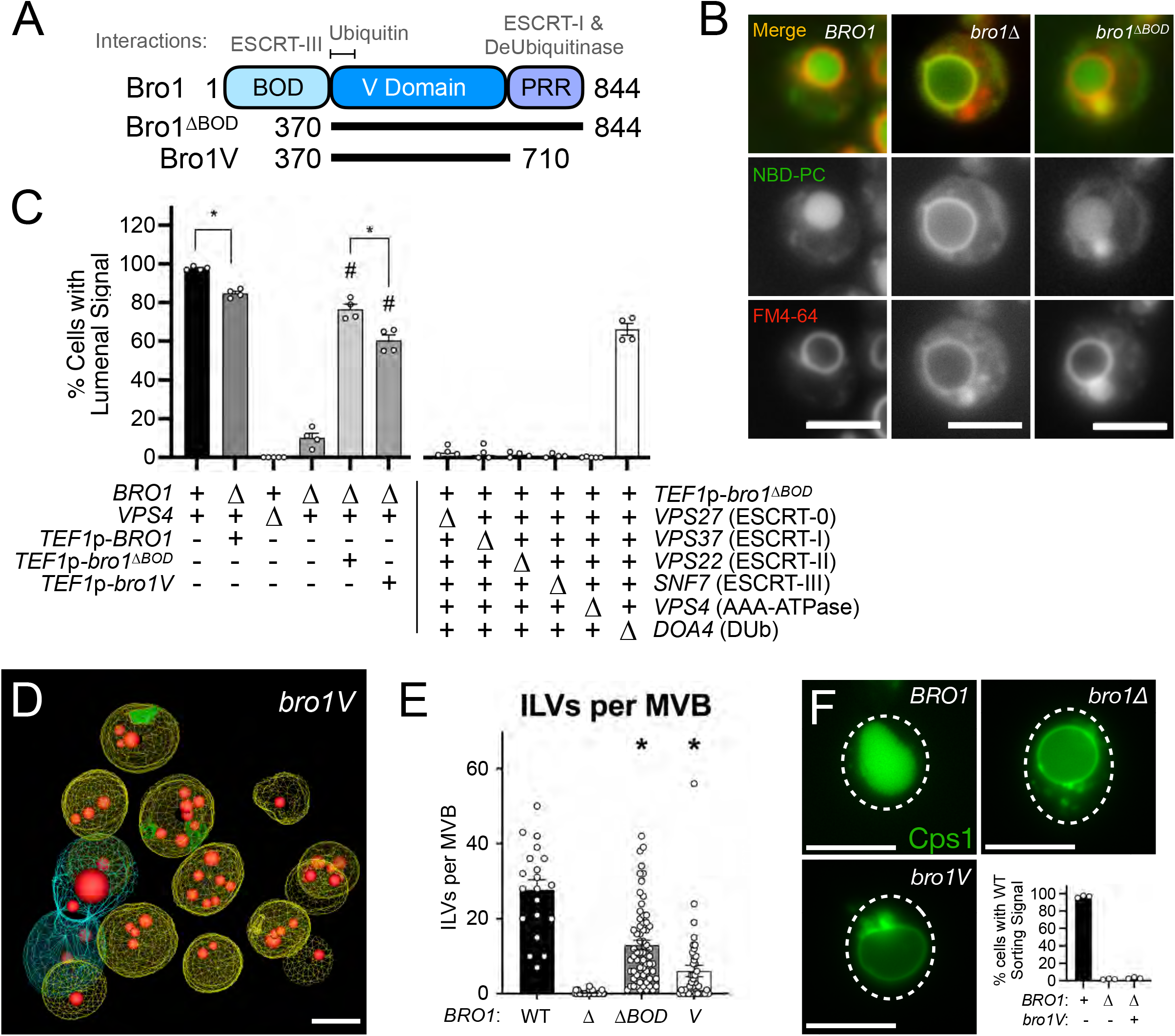
Bro1ΔBOD sorting of NBD-PC into the vacuolar lumen is dependent on Vps4/ESCRT machinery. **(A)** Domain cartoon of Bro1 (amino acids 1-844) with interacting factors annotated. **(B)** Sample micrographs of WT (SEY6210), *bro1*Δ (GOY65), and *bro1*^Δ*BOD*^ (*bro1*Δ::*TEFL1*p-*bro1*^Δ*BOD*^ CTY2) cells co-stained with NBD-PC and FM4-64 revealing endpoints of the observed phenotypes. Scale = 5μm. **(C)** NBD-PC and FM4-64 stained WT (SEY6210), *TEF1*p-*BRO1* (CTY1), *vps4*Δ (MBY3), *bro1*Δ *(GOY65), TEF1*p-*bro1*^Δ*BOD*^ (CTY2), *TEF1*p-*bro1V*(CTY4), *bro1*^Δ*BOD*^ *vps4*Δ (CTY5), *bro1*^Δ*BOD*^ *vps27*Δ (CTY29), *bro1*^Δ*BOD*^ *vps37*Δ (CTY21), *bro1*^Δ*BOD*^ *vps22*Δ (CTY24), *bro1*^Δ*BOD*^ *snf7*Δ (CTY12), *bro1*^Δ*BOD*^ *vps4*Δ(CTY5), and *bro1*^Δ*BOD*^ *doa4*Δ(CTY13) cells were analyzed by live cell fluorescence microscopy and quantified for the frequency of cells able to support NBD-PC trafficking to the vacuolar lumen. Error bars indicate S.E.M. * indicates statistically significant differences. # indicates statistically significant difference compared to *bro1*Δ. Percentage of cells with WT sorting signal was quantified and calculated from at least 94 cells from four independent experiments performed on four different days. White dashed lines indicate cell boundaries. Scale bar = 5μm. **(D, E)** *bro1V* (CTY4) were analyzed by electron tomography and ILVs per MVB were quantified. 3D reconstructions of the tomogram are shown in **(D)**. The limiting membrane of normal-like MVBs are labeled yellow, while the limiting membrane of tubular/aberrant MVBs are shown in different colors. ILVs are highlighted in red, A minimum of 13 MVBs from at least 10 cells were quantified. Scale bar = 100nm. **(F)** WT (SEY6210), *bro1*Δ (GOY65), or *bro1V*(*bro1*Δ::*TEFL1*p-*bro1V*; CTY4) cells were transformed with the GFP-CPS plasmid to assess MVB sorting using live cell fluorescence microscopy. Percentage of cells with WT sorting signal was quantified and calculated from at least 156 cells from three independent experiments performed on three different days from two different transformations. White dashed lines indicate cell boundaries. Scale bar = 5μm.

Bro1^ΔBOD^ contains both the V domain as well as a C-terminal proline rich domain (PRR) that mediates interaction with the Ub isopeptidase Doa4 (Buysse et al., 2020). We found that Bro1^ΔBOD^-mediated NBD-PC sorting was not dependent on Doa4 and that Bro1 V domain alone supports NBD-PC sorting into the vacuole lumen (Figure 3C), demonstrating that PRR and its associations are not required for this activity. This finding was supported by EM tomography analysis wherein Bro1 V domain alone (*bro1V*) was able to support some ILV formation (Figure 3D, Movie S5), albeit at levels less than WT or *bro1*^Δ*BOD*^ cells (Figure 3E). However, the size of the ILVs in *bro1V* cells were indistinguishable from those in WT or *bro1*^Δ*BOD*^ cells (Figure S2C). While Bro1V alone is able to support ILV formation, Bro1V was unable to support MVB cargo sorting (Figure 3F, Figure S2D), as observed with *bro1*^Δ*BOD*^ cells. In total, these results implicate the V domain in promoting Bro1/Vps4/ESCRT-III-driven ILV formation.

### Bro1 V domain stimulates Vps4 in vitro

The ability of Bro1^ΔBOD^ and, in particular, the Bro1 V domain alone to support ILV formation without promoting efficient MVB protein cargo sorting suggested that the V domain might activate ESCRT function at the level of ESCRT-III/Vps4 membrane remodeling and scission activity *in vivo* in a manner that could be uncoupled from efficient cargo packaging. Thus, we searched for how the V domain might directly stimulate the vesicle scission machinery by examining effects on Vps4 activity. Bro1 interacts with Vps4 (Vajjhala et al., 2007), however the mode and significance of this association have not been determined. Binding studies using a series of recombinant GST-tagged Bro1 protein fragments and His_6_-tagged Vps4 protein fragments showed that the Bro1V binds the N-terminal Vps4 MIT domain (Figure 4A,B). Two distinct surfaces of the MIT domain mediate interactions with MIM1 and MIM2 elements in ESCRT-III subunits (Kieffer et al., 2008; Obita et al., 2007; Stuchell-Brereton et al., 2007), however mutations (I18D, MIM2; L64D, MIM1) disrupting these modes of association did not impair MIT-V domain association. These results suggest that Bro1V binds Vps4 MIT domain in a manner distinct from MIM1 and MIM2.

**FIGURE 4.**
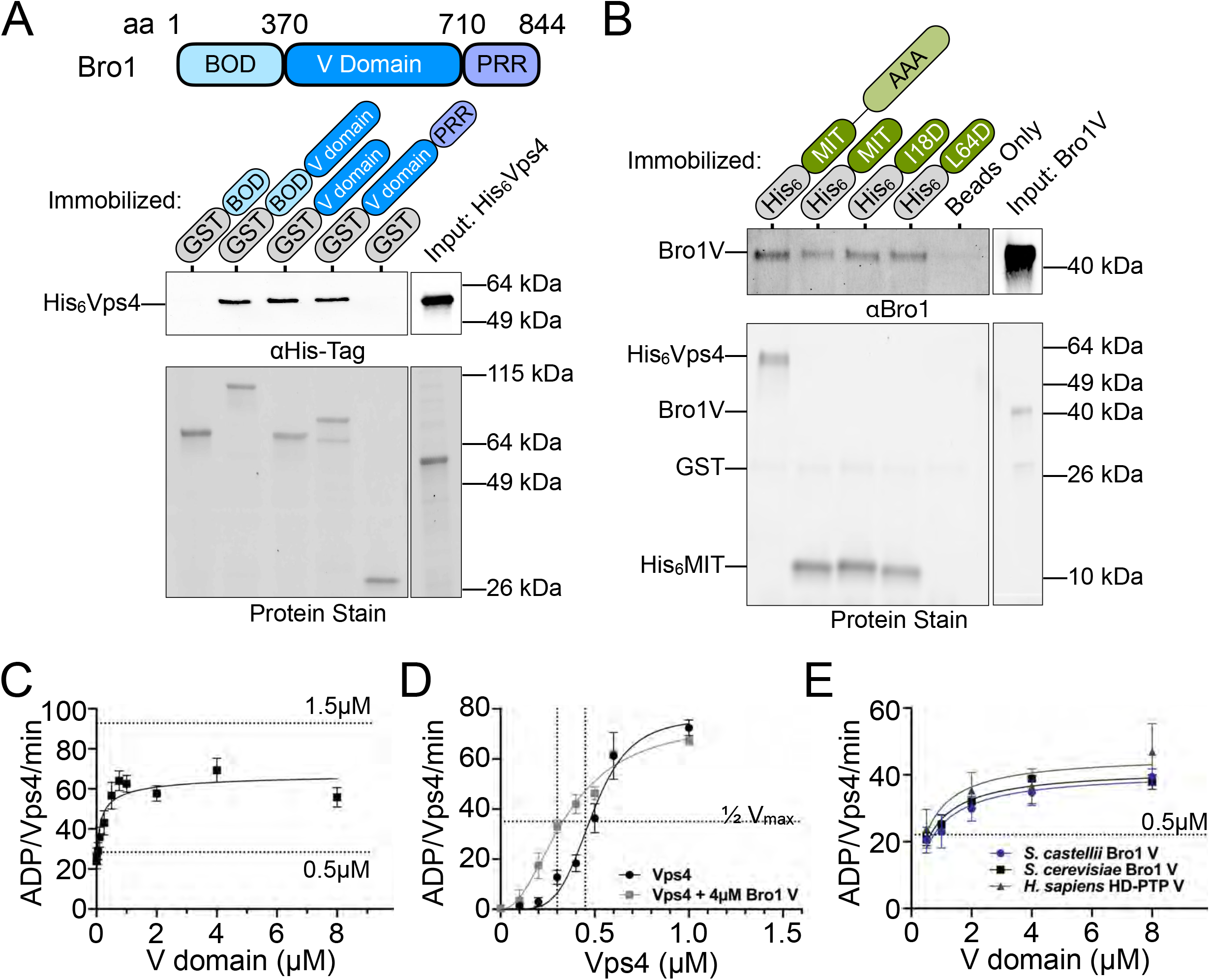
V domain stimulates Vps4 ATPase activity *in vitro*. **(A)** GST-Bro1 fragments and GST bound beads were incubated with His_6_-Vps4. Bound material was visualized by both EZBiolab Instant-Band Protein Stain and immunoblotting with penta-His antibody. **(B)** His_6_-Vps4, His_6_-MIT and Ni-NTA beads were incubated with Bro1V. Bound material was visualized by both EZBiolab Instant-Band Protein Stain and immunoblotting with anti-Bro1 antiserum. **(C)** Vps4 specific activity with titration of Bro1V (10nM-8μM). Vps4 (0.5μM) ATPase assays were conducted using the indicated conditions and resolved by thin layer chromatography for quantitation and calculation of hydrolysis rates. Dashed lines indicate Vps4 specific activity for 0.5μM or 1.5μM Vps4 alone, as indicated. Bro1V alone did not exhibit measurable ATP hydrolysis. **(D)** Vps4 titrations were performed with or without 4μM Bro1V. Vps4 specific activity is presented. The vertical dotted line indicates the Vps4 apparent K_m_ +/- Bro1V. **(E)** V domains of *S. cerevisiae* Bro1, *S. castellii* Bro1, and *H. sapiens* HD-PTP (0.5-4μM) were titrated against 0.5μM Vps4. Specific activity of Vps4 is expressed as ADP generated per Vps4 molecule per minute. Error bars indicate S.E.M.

The possibility that Bro1V domain modulates Vps4 activity was assessed using ATPase assays with purified recombinant proteins (Babst et al., 1997). Vps4 exhibits concentration dependent increases in specific activity due to Vps4 oligomer formation with maximal activity observed with 1.5μM Vps4 (Azmi et al., 2006; Babst et al., 1998; Davies et al., 2010). Titration of Bro1V into ATPase assays with a submaximal concentration of Vps4 (0.5μM) showed a concentration dependent stimulation of activity with a maximum of 2.6-fold over basal activity (Figure 4C). Stimulated activity of 0.5μM Vps4 did not surpass the inherent maximal Vps4 specific activity observed with 1.5μM Vps4, and Bro1V was unable to stimulate 1.5μM Vps4 specific activity (data not shown). These observations suggested that Bro1V stimulates Vps4 oligomerization without further enhancing activity of the Vps4 oligomer. To examine this directly, Vps4 titration was performed in the presence of 4μM Bro1V. While Bro1V enhanced Vps4 specific activity up to 0.6μM Vps4, Vps4 concentrations exhibiting near maximal specific activity (0.6-1.0μM Vps4 in this particular series of experiments) were not further stimulated by Bro1V (Figure 4D). Nonlinear regression indicated that Bro1V addition reduced the Vps4 apparent K_m_ from 0.45μM to 0.30μM, supporting enhanced oligomerization as the mechanism by which Bro1 stimulates Vps4. Furthermore, V domains from *S. castellii* Bro1 (residues 370-709) and human HD-PTP (residues 364-695) stimulated Vps4 ATPase activity in a similar dose-dependent manner (Figure 4E). Together, these results reveal that Bro1 V domain enhances Vps4 ATPase activity directly, and suggest this stimulatory activity is conserved among homologs.

### Generation of Bro1 V domain mutants defective for Vps4 stimulation

We next wanted to determine the functional significance of the stimulatory effect of Bro1V on Vps4 activity. In addition, we wanted to determine whether this effect was due solely to simple binding of Bro1V to Vps4 or whether additional biochemical processes could be observed that were responsible for stimulating Vps4. Therefore, we mutated select residues that were conserved between Bro1 and HD-PTP with the goal of generating Bro1 mutants specifically altered in in their ability to stimulate Vps4 activity but retained their ability to bind Vps4 (Figure 5, Figure S3A). Vps4 stimulation was measured using ATPase reactions containing WT or mutant forms of Bro1V. Six of the Bro1V mutants tested stimulated Vps4 activity like WT Bro1V, whereas four mutants — **Bro1V^M4^**: V505A, H508A, I512A; **Bro1V^M8^**: E686A, L691D; **Bro1V^M9^**: T587D; and **Bro1V^M10^**: K481D — did not stimulate Vps4 ATPase activity (Figure 5B) but retained their ability to bind Vps4 (Figure 5C, Figure S3A). Indeed, ability of these mutants to associate with Vps4, in some cases better than WT, led us to perform titration experiments to assess the impact on Vps4 ATPase activity. While the Bro1V mutants were unable to stimulate Vps4 at higher concentrations (>1μM), these mutants could stimulate Vps4 ATPase activity at lower concentrations (≤1μM; Figure 5D). This profile contrasted with the behavior of WT Bro1V that continued to stimulate Vps4 ATPase activity even at the highest concentrations tested. Together these analyses suggest that Bro1V stimulation of Vps4 relies on determinants beyond mere binding and provided tools to determine whether Bro1V-stimulated Vps4 activity was important *in vivo* for MVB sorting and biogenesis.

**FIGURE 5.**
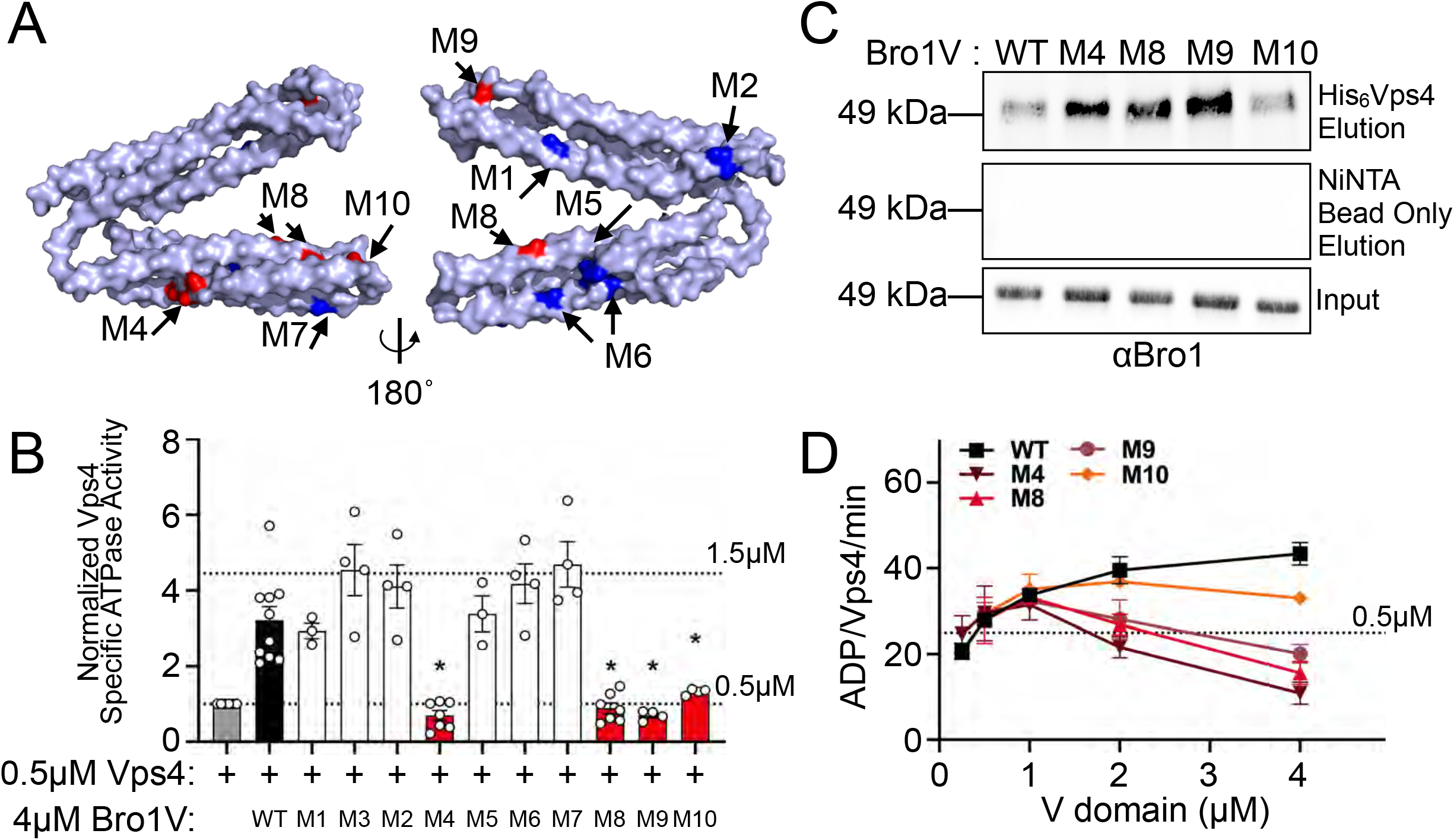
Bro1 V domain mutations disrupt Vps4 stimulation *in vitro* without disrupting binding. **(A)** Model of the V domain based on the *S. castellii* Bro1 V domain crystal structure (Protein Data Bank ID: 4JIO, chain A) with amino acid substitution mutations impacting V domain stimulation of Vps4 indicated in red. Conserved residues that when mutated did not impact Vps4 stimulation *in vitro* are indicated in blue. See Supplemental Table 4 for individual mutations. **(B)** Stimulation of Vps4 ATPase activity (0.5μM) by 4μM Bro1V(370-709) and Bro1V mutants represented as normalized Vps4 ATPase activity of at least three experiments done in duplicate. Error bars indicate S.D. * indicates statistically significant differences compared to WT. **(C)** Immobilized His_6_-Vps4 or Ni-NTA beads alone were incubated with Bro1V, Bro1V(M4), Bro1V(M8), Bro1V(M9), and Bro1V(M10). Bound material was visualized by immunoblotting with anti-Bro1 antiserum. **(D)** Vps4 (0.5μM) ATPase activities with titration of Bro1V, Bro1V(M4), Bro1V(M8), Bro1V(M9), and Bro1V(M10) (0.25-5μM). Vps4 specific activity is expressed as ADP generated per Vps4 molecule per minute. Error bars indicate S.E.M.

### Altered regulation of Vps4 activity by Bro1V perturbs ILV formation in vivo

MVB sorting was examined in *bro1*Δ cells carrying full-length alleles of *BRO1* containing mutations in the V domain that altered Vps4 stimulation (Figure 6A). Three distinct model MVB cargoes were utilized for this analysis: GFP-Cps1 for its strict dependence upon Ub modification for proper sorting, the chimera Ub-GFP-Cps1 for its lack of dependence upon exogenous Ub modification, and Sna3-GFP as a cargo that has been shown to access the MVB pathway both directly and via association with other ubiquitylated cargoes (Katzmann et al., 2001; Katzmann et al., 2003; MacDonald et al., 2012b; Odorizzi et al., 1998; Oestreich et al., 2006; Reggiori and Pelham, 2001). Although all Bro1 mutants were expressed to levels similar to WT Bro1, none of the mutant strains (*bro1^M4^, bro1^M8^, bro1^M9^*, and *bro1^M10^*) mediated MVB sorting of any cargo (Figure 6A-C, Figure S3B). The correlation between these defects in MVB cargo sorting and altered Vps4 stimulation observed with these mutants at higher concentrations support the conclusion that proper Bro1 V domain stimulation of Vps4 is required for ESCRT-driven MVB cargo sorting *in vivo*.

**FIGURE 6.**
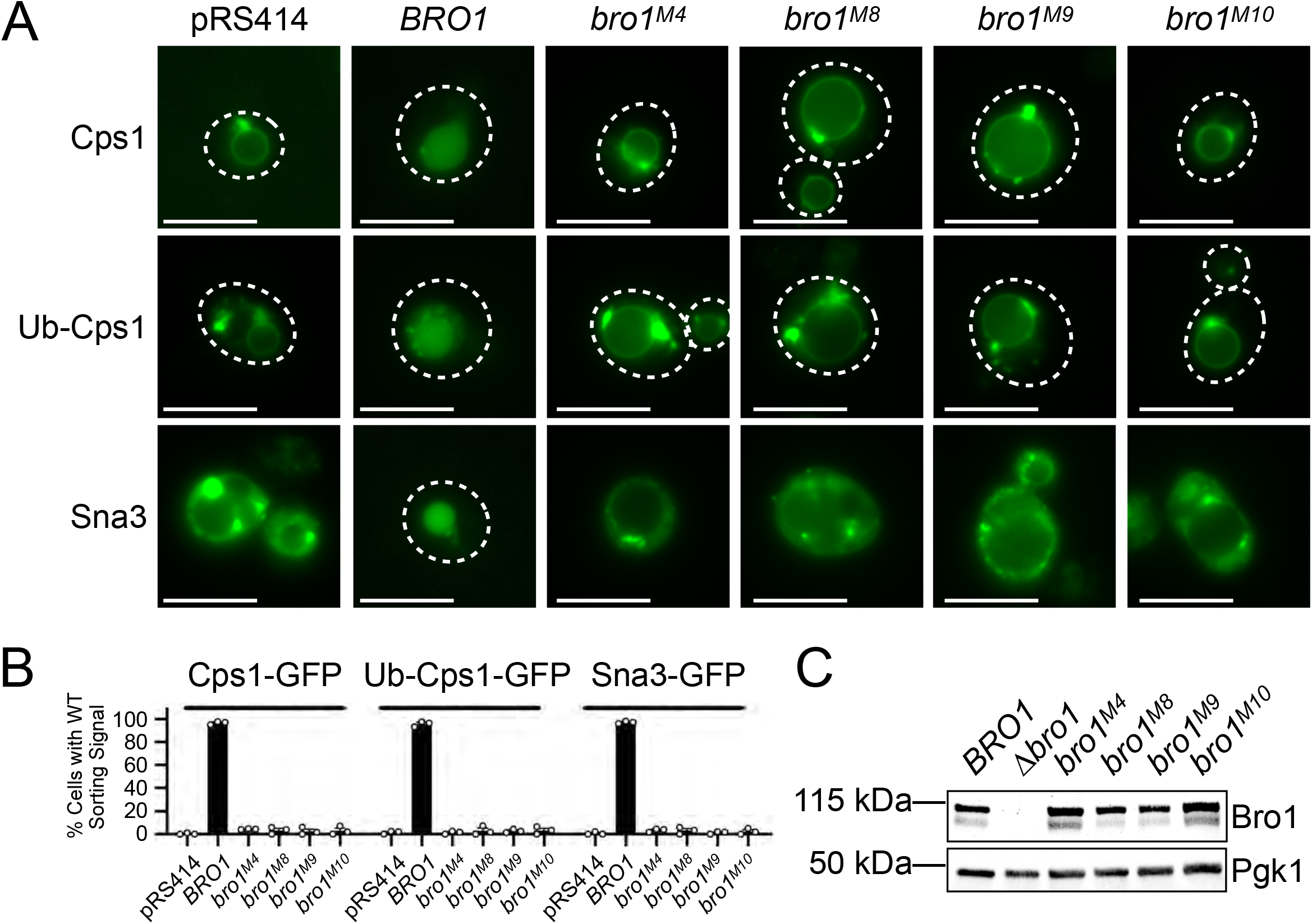
Bro1 V domain mutations disrupting Vps4 stimulation *in vitro* disrupt MVB sorting *in vivo*. **(A)** *bro1*Δ (GOY65) cells were transformed with empty plasmid (pRS414) or plasmids with the *BRO1* promoter and *BRO1, hro1(M4), bro1(M8), bro1(M9)*, or *bro1(M10)*. Localizations of model MVB cargo GFP-Cps1, Ub-GFP-Cps1 or Sna3-GFP were determined using live cell fluorescence microscopy to assess MVB sorting in these mutant contexts. White dashed lines indicate cell boundaries. Scale bar = 5μm. (**B)** Percentage of cells with WT sorting signal was quantified and calculated from at least 100 cells from three independent experiments performed on three different days from three different transformations. **(C)** Representative immunoblots showing mutant protein expression levels, probing against Bro1 and PGK1 as a loading control, using lysates of GOY65 transformed with empty plasmid (pRS414) or plasmids with the *BRO1* promoter and *BRO1, bro1(M4), bro1(M8), bro1(M9)*, or *bro1(M10)*.

Dual axis electron tomography was used to examine how loss of normal Bro1-stimulated Vps4 activity altered the formation of ILVs (Figure 7A). ILVs of mutant *bro1^M8^cells* were the same size as those in WT cells. However, *bro1^M8^* cells had roughly twice as many ILV budding intermediates per MVB compared to WT cells as well as reduction in the surface area of the budding profiles (Figure 7B, Movie S4). These observations suggest that proper stimulation of Vps4 by Bro1V contributes to ILV formation at the level of bud expansion.

**FIGURE 7.**
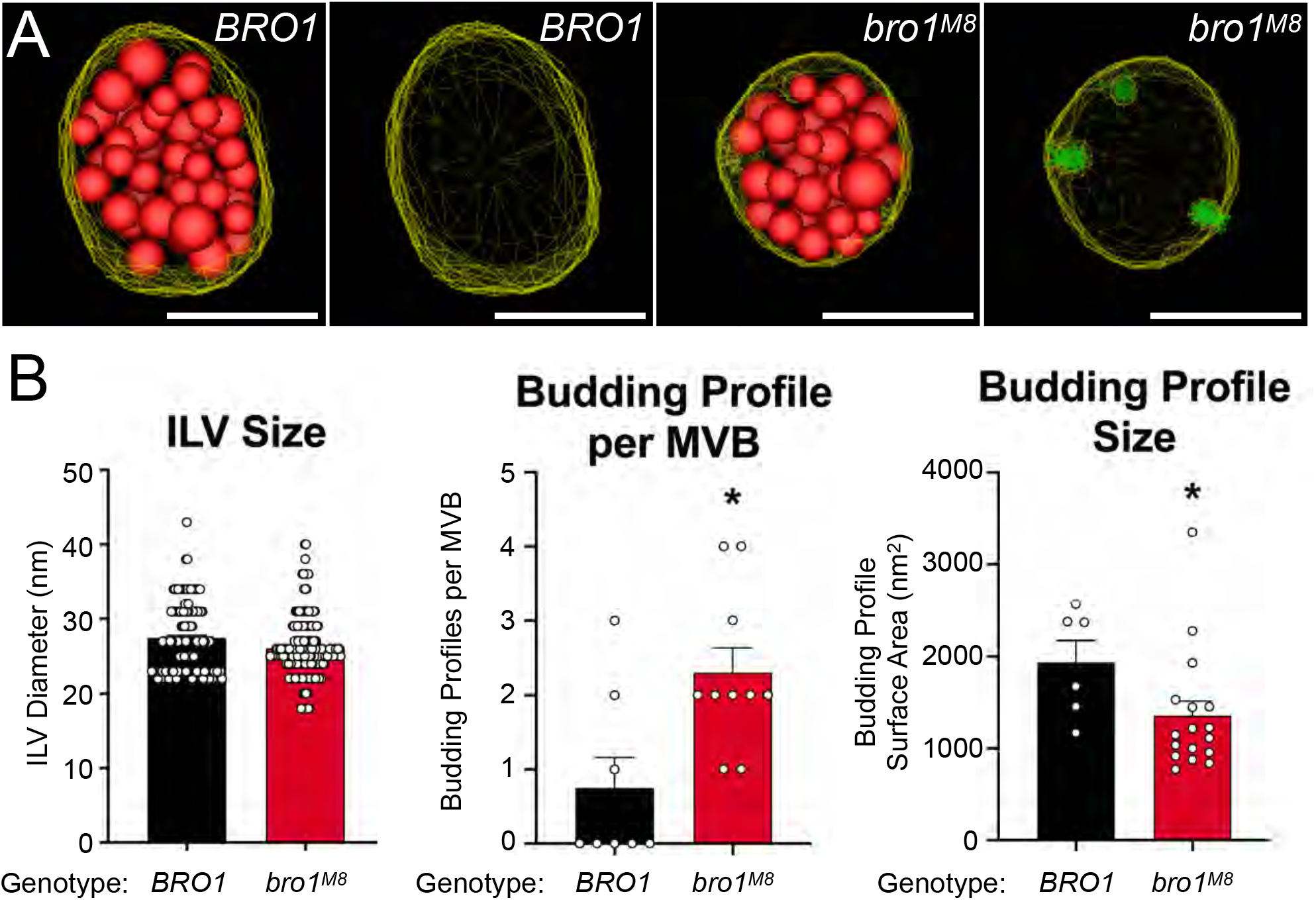
Bro1(M8) perturbs ILV formation *in vivo*. **(A)** Three-dimensional models reconstructed from 200-nm-thick section electron tomograms of *bro1*Δ (GOY65) with *BRO1* or *bro1(M8)* plasmids. The limiting membrane of the MVB is labeled yellow, the ILVs are highlighted in red and the budding intermediates are colored in green. Scale bar = 100nm. **(B)** Quantification of electron tomograms of GOY65 with either *BRO1* or *bro1(M8)* plotting ILV size, Budding Profile Size, and Budding Profiles per MVB. A minimum of 9 MVBs and 208 ILVs from at least 10 cells were measured. Error bars indicate S.E.M. * indicates statistically significant differences compared to WT.

We also tested whether Bro1 stimulation of Vps4 regulation affected the steady-state membrane association of ESCRT-III, which is a measure of the polymeric state of ESCRT-III subunits. Subcellular fractionation of *bro1^M4^* and *bro1^M8^* cells showed WT levels of the ESCRT-III subunit Snf7 on membranes in contrast to *bro1*Δ cells that showed more soluble Snf7 (Figure 8). These findings suggest that ESCRT-III polymerization and disassembly/recycling were normal in *bro1^M4^* and *bro1^M8^* cells, implying that Bro1V-mediated Vps4 stimulation exerts its function at a different biochemical step.

**FIGURE 8.**
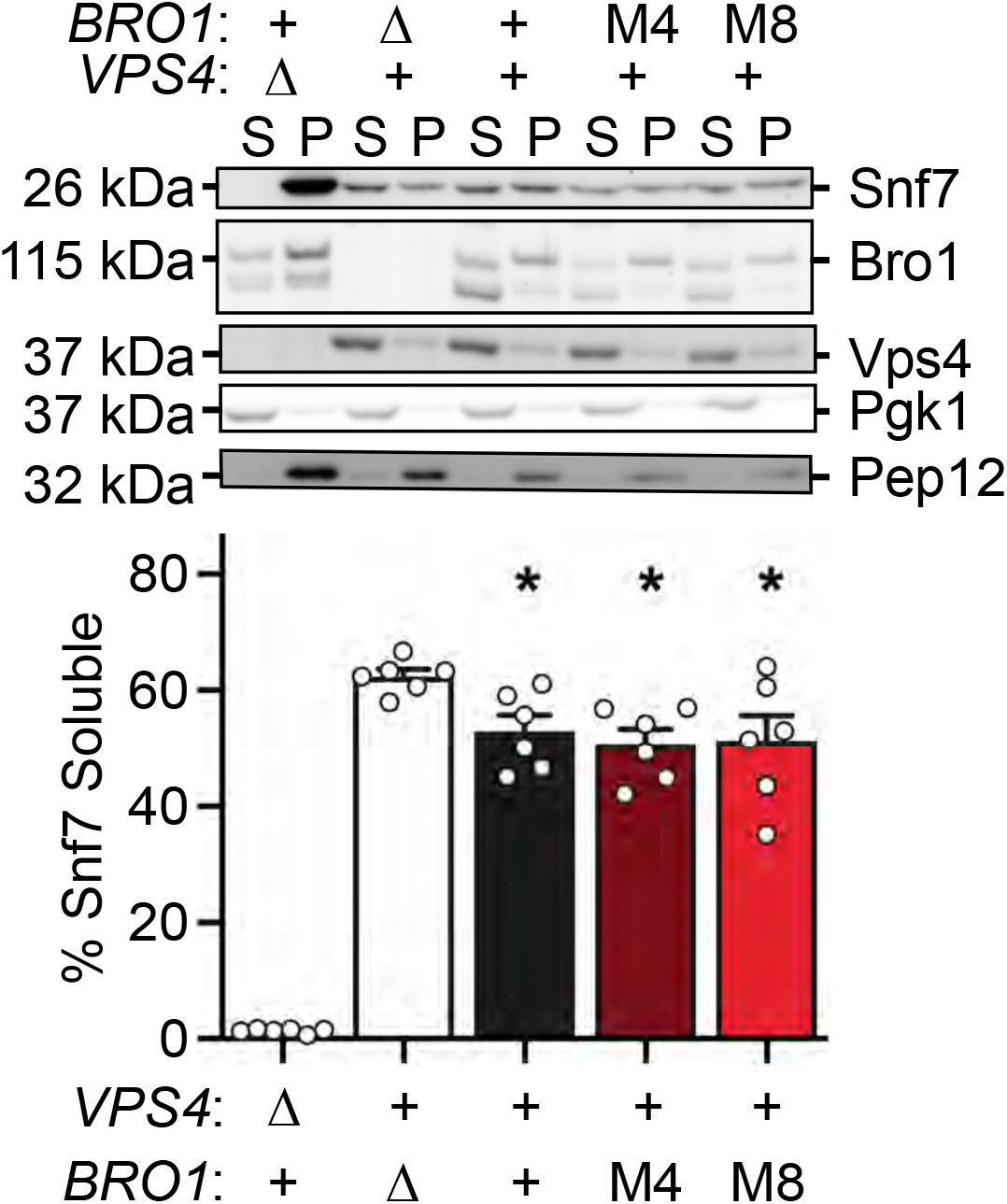
Bro1(M4) and Bro1(M8) display WT steady state membrane-associated ESCRT-III *in vivo*. Subcellular fractionation was performed in *bro1*Δ (GOY65) cells transformed with an empty vector or *BRO1, bro1(M4)*, or *bro1(M8)* plasmids. *vps4*Δ (MBY3) cells were used as a control highlighting the distribution upon complete loss of Vps4 function. Representative immunoblots indicating fractionation of Snf7, Bro1, and Vps4 are shown. Pep12 and Pgk1 were used as membrane and soluble markers respectively. Quantification represents six experiments; Error bars indicate S.E.M. * indicates statistically significant difference compared to *bro1*Δ.

### Ub binding enhances Bro1 V domain stimulation of Vps4 activity in vitro

Bro1 binds Ub via a conserved motif at the amino-terminal portion of the V domain (Figure S4A, S4B) (Pashkova et al., 2013). While Bro1 has been implicated as a Ub-sorting receptor (Pashkova et al., 2013), whether Ub-binding impacts other aspects of MVB biogenesis is unclear. Therefore, we examined if Ub-binding to the V domain impacts its stimulation of Vps4 (Figure 9A). While Ub does not directly stimulate Vps4 *in vitro*, addition of Ub to ATPase reactions increased Bro1V-stimulation of Vps4 activity using Bro1V from both *S. cerevisiae* and the related yeast *S. castellii*, for which the crystal structure is known (Pashkova et al., 2013). In contrast, addition of Ub to mutant Bro1V^ΔUBD^ (I377R) in which Ub-binding was ablated had no effect on Vps4 stimulation. These results indicate a conserved mode of Ub-dependent regulation of the V domain’s impact on Vps4.

**Figure 9.**
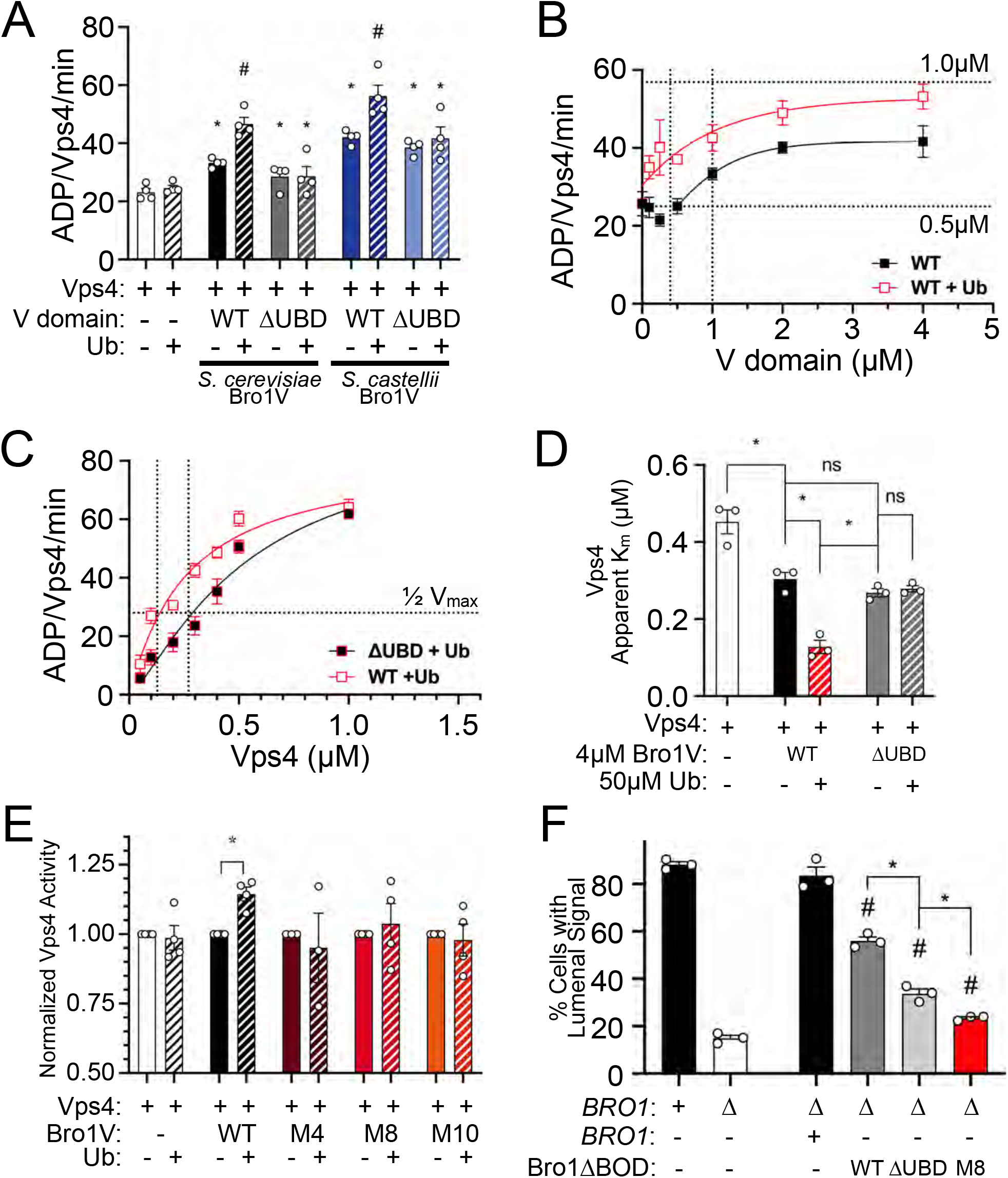
Ubiquitin potentiates V domain stimulation of Vps4 ATPase activity. **(A)** Vps4 (0.5μM) ATPase specific activity in the presence of 1μM *S. cerevisiae* Bro1V, *S. cerevisiae* Bro1V^ΔUBD^ (I377R), *S. castellii* Bro1V(370-708) and *S. castellii* Bro1V^ΔUBD^ (I377R) +/- 50μM mono-Ub. Error bars indicate S.E.M. * indicates statistically significant difference compared to Vps4 alone. # indicates statistically significant difference compared to Vps4 + Bro1V - Ub. **(B)** Bro1V titration performed in the presence of 0.5μM Vps4 with or without 50μM Ub. The vertical dotted line indicates the Bro1V concentration generating half-maximal stimulation within each context. **(C)** Vps4 titration (0.05-1.0 μM) in the presence of 4μM Bro1V WT or Ub binding mutant (L386R) and 50μM Ub. Vps4 specific activity (ADP generated per Vps4 molecule per minute) is presented. Error bars indicate mean +/- S.E.M. Vertical dotted lines indicate the Vps4 apparent K_m_ in each context. **(D)** The Vps4 apparent K_m_ of Vps4 alone, Vps4 + 4μM Bro1V, Vps4 + 4μM Bro1V + 50μM Ub, Vps4 + 4μM Bro1V^ΔUBD^ (L386R), and Vps4 + 4μM Bro1V^ΔUBD^ + 50μM Ub, was determined from Vps4 titration experiments in Figure 4D, Figure 9C, and Figure S4C. * indicates statistically significant difference. **(E)** Vps4 ATPase activity (0.5μM) in the presence of 4μM *S. cerevisiae* Bro1V, Bro1V^M4^, Bro1V^M8^ or Bro1V^M10^ without or with 50μM of mono-Ub. Vps4 specific activity with Ub addition is normalized to activity without Ub. Error bars indicate S.E.M. * indicates statistically significant difference +/- Ub addition. **(F)** NBD-PC and FM4-64 stained WT (SEY6210), *bro1*Δ (GOY65) and GOY65 reexpressing Bro1 or over-expressing Bro1^ΔBOD^, Bro1^ΔBOD,ΔUBD^ (I377R, L386R), and Bro1^ΔBOD,M8^ were analyzed by live cell fluorescence microscopy and quantified for the frequency of cells with NBD-PC in the vacuolar lumen. Error bars indicate S.E.M. * indicates statistically significant difference. # indicates statistically significant difference compared to *bro1*Δ.

We next examined how Ub impacts the stimulation of Vps4 by Bro1V using titration experiments in presence or absence of Ub (Figure 9B). Ub increased the potency of Bro1V wherein the Bro1V concentration yielding half-maximal stimulation of Vps4 was reduced from 1.0μM to 0.4μM. We also examined the impact of Ub-bound Bro1V on Vps4 oligomer assembly and/or activity of the assembled Vps4 oligomer. In these experiments, Vps4 titration was performed in the presence of Ub and either WT Bro1V or a Bro1V mutant unable to bind Ub, Bro1V^ΔUBD^ (L386R). Whereas both WT Bro1V and Bro1V^ΔUBD^ similarly reduced the apparent K_m_ of Vps4, addition of Ub to WT Bro1V, but not to Bro1V^ΔUBD^, further reduced the Vps4 apparent K_m_ (Figure 9C, 9D, S4C). These results suggest that Ub-binding alters V domain conformation to enhance its association with Vps4 and promote Vps4 oligomerization without impacting activity of the assembled Vps4 oligomer.

We next tested whether the Bro1V mutants defective for Vps4 stimulation retained their capacity to bind Ub as well as undergo Ub-mediated enhancement of Vps4 stimulation. The Bro1V mutants impacting Vps4 stimulation are not confined to a single surface nor do they disrupt association with Vps4, suggesting that these mutations alter the conformation or dynamics of the V domain, which in turn is used to stimulate Vps4. Whereas Bro1V(I377R) failed to bind Ub, Bro1V^M4^, Bro1V^M8^, Bro1V^M10^, and WT Bro1V all bound Ub (Figure S4B, S4D). The Bro1V^M4^, Bro1V^M8^, and Bro1V^M10^ mutants defective in Vps4 stimulation also showed none or diminished Ub-mediated enhancement of Vps4 stimulation (Figure 9E). Together these results demonstrate that Ub enhances V domain activity in a manner dependent on V domain Ub-binding and V domain stimulation of Vps4.

### Ub promotes ILV formation via the Bro1/Vps4/ESCRT-III axis

Bro1 has been proposed to work early in the MVB sorting process as a receptor for Ub-cargo. Defects in cargo sorting by Bro1 mutants defective in Ub-binding are revealed when a parallel cargo recognition pathway mediated by ESCRT-0 is compromised (Pashkova et al., 2013). Because this function is upstream of the Vps4/ESCRT-III ILV formation axis, it was not clear whether Ub-binding to Bro1V might play a functional role in downstream events. Since Bro1 lacking its BOD drives ILV formation without efficient cargo sorting, we examined whether Ub-binding might contribute to this latter activity by following NBD-PC sorting. Cells expressing Bro1^ΔBOD^ defective for Ub binding (ΔUBD: I377R, L386R) showed reduced levels of NBD-PC sorting in comparison with WT indicating that Ub may play a role in ILV formation itself (Figure 9F); similar results were observed with deletion of the Ub-binding residues (aa388-844, Figure S2B). In addition, mutant Bro1^ΔBOD,M8^ which contains a V domain unable to properly stimulate Vps4 showed a further reduction in NBD-PC sorting, barely above sorting observed in *hro1Δ* cells. These degrees of NBD-PC sorting with Bro1^ΔBOD,ΔUBD^ and Bro1^ΔBOD,M8^ correlate with Bro1V^ΔUBD^ exhibiting basal but not Ub-enhanced stimulation of Vps4 *in vitro* (Figure 9A). These results indicate that 1) V domain stimulation of Vps4 contributes to Bro1^ΔBOD^ ILV formation, and 2) Bro1 Ub-binding enhances ILV formation via the Bro1/Vps4/ESCRT-III axis.

## Discussion

Here we reveal a new function of Bro1 in MVB biogenesis that is mediated by its V domain binding to and stimulating Vps4 (Figure 10). This stimulation is further enhanced by Ub-binding, indicating that Bro1 interaction with Ub-cargo may regulate Vps4 activity via this biochemical mechanism. Moreover, this mode of Vps4 regulation is important for proper ILV formation and cargo sorting into ILVs. These data suggest Bro1 plays a role in “licensing” ESCRT-III and Vps4 to drive membrane remodeling in concert with cargo transfer into the ILV. The idea that Bro1 couples cargo sorting with ILV formation is underscored by the observations here and elsewhere that disconnecting Bro1 from some of its interactions separates the process of ILV formation with the process of efficiently sorting cargos into those ILVs. We propose that the central Bro1 activity of the V domain stimulating Vps4/ESCRT-III-driven ILV formation is coordinated with BOD and PRR activities to properly time ILV budding to enable normal MVB biogenesis.

**Figure 10.**
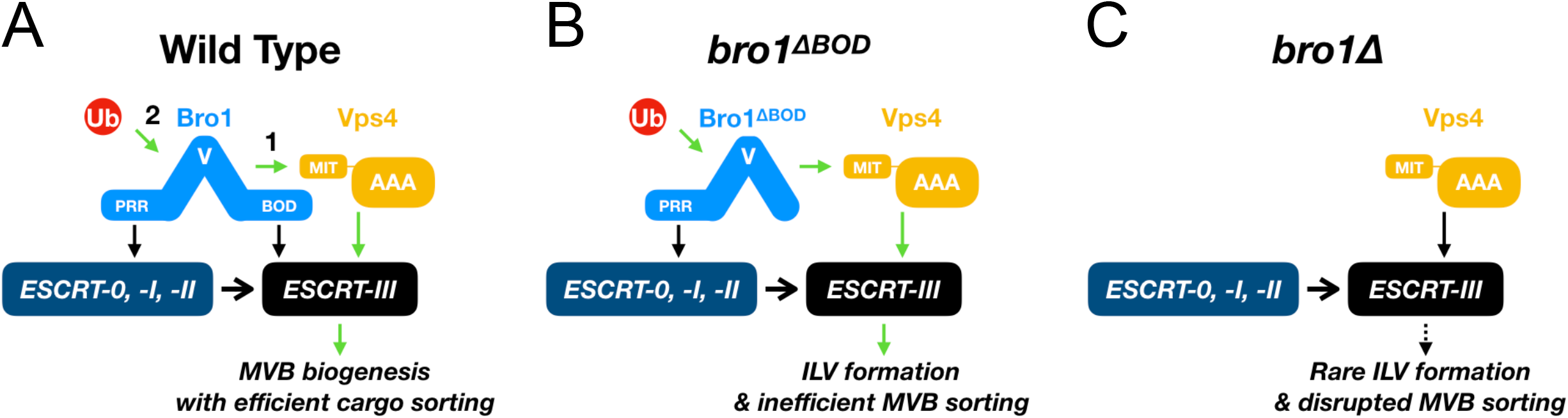
Model of Bro1 “licensing” ESCRT-III membrane remodeling. **(A)** Bro1 interacts with ESCRTs via interactions between Proline Rich Region (PRR) and early ESCRTs (−0, -I & - II) as well as Bro1 Domain (BOD) and ESCRT-III subunit Snf7. **1)** Bro1 V domain binds Vps4 MIT domain, and promotes Vps4 activity. This stimulation promotes ILV formation and is required to coordinate cargo sorting and ILV formation. **2)** Stimulation of Vps4 by Bro1 V domain is further enhanced by Ub, though the exact source of Ub remains unclear. **(B)** In the context of *bro1*^Δ*BOD*^, V domain stimulation of Vps4 promotes ILV formation without efficient cargo sorting. **(C)** Vps4 and ESCRT-III cannot efficiently generate ILVs in the absence of Bro1 and MVB sorting is disrupted.

Bro1 is one of several proteins that regulate Vps4 activity. Vta1 and ESCRT-III subunits also stimulate Vps4 activity (Azmi et al., 2006; Azmi et al., 2008; Merrill and Hanson, 2010; Shim et al., 2008), but do so by connecting differently to Vps4. We found that the Bro1 V domain binds to the Vps4 MIT domain and increases Vps4 specific activity by reducing its apparent Km, suggesting that Bro1V facilitates Vps4 oligomerization. The Vps4 MIT domain also binds several ESCRT-III subunits along two distinct surfaces that mediate association with MIM1 and MIM2 motifs found in Vps2 or Vps24 and Vps20 or Snf7, respectively (Kieffer et al., 2008; Obita et al., 2007; Stuchell-Brereton et al., 2007). Mutations that disrupt MIM1 or MIM2 binding did not alter the ability of Bro1V to bind the MIT domain, suggesting that Bro1V binds by a distinct mode. Yet it remains to be determined if Bro1V binding diminishes or enhances the ability of Vps4 to bind to MIM1 or MIM2-containing ESCRT-III subunits. Vps4 recruitment to the site of MVB sorting occurs in two phases: an early, minimal recruitment of Vps4 associated with stable ESCRT-III membrane association followed by a later recruitment of additional Vps4 implicated in ILV formation and eventual ESCRT-III disassembly (Adell et al., 2017). Our *in vitro* activity analyses indicate that Bro1 facilitates Vps4 oligomerization, as Bro1V reduces the apparent K_m_ of Vps4 itself. Yet, exactly how Bro1V regulation of Vps4 activity impacts the dynamics and function of the Vps4 substrate polymer ESCRT-III remains to be determined. Overall, the level of polymerized membrane-associated ESCRT-III remains unperturbed in cells where Bro1 is unable to properly stimulate Vps4 ATPase activity. Moreover, the size of the ILVs themselves, an indicator of the fidelity of ESCRT-III mediated scission, is unaltered as well. Together, these data suggest this role of Bro1V regulation is transient, precise and distinct from Vta1/ESCRT-III stimulation of Vps4 implicated in ILV scission and ESCRT-III disassembly. Bro1 mutants with altered Vps4 regulation did exhibit reduced budding profile size, suggesting Bro1 stimulation of Vps4 contributes to ILV bud expansion. Bud expansion controlled by Vps4 should be coordinated with entry of cargo as well as deubiquitination of cargo just prior to its entry into ILVs, and the biochemical activities now described for Bro1 position it well for such a role. While future studies are required to expand our understanding of Bro1’s role in the Vps4/ESCRT-III ILV formation axis, our studies unequivocally demonstrate Bro1 stimulation of Vps4 makes unique contributions to MVB biogenesis.

Analyses *in vitro* and *in vivo* indicate that V domain mutations we characterized retain partial Vps4 stimulation activity. These mutants retain interaction with Vps4 and Ub and stimulation of Vps4 ATPase activity at lower concentrations, however these mutants are defective for MVB cargo sorting *in vivo* and stimulation of Vps4 ATPase activity at higher concentrations in vitro. We conclude Bro1V stimulation of Vps4 observed *in vitro* under high concentrations is the activity relevant for Bro1 function in MVB sorting *in vivo*. These mutations map to both arms of the V domain, suggesting they impart subtle conformational changes in V domain that maintain association with Vps4 but disrupt stimulation. Further structural studies are underway to elucidate this mechanism.

Our results also suggest that Vps4 responds to Ub through Bro1. Titration experiments indicate that Ub-binding to the V domain enhances its ability to stimulate Vps4 via facilitating Vps4 oligomerization. Previous studies demonstrated a role for Ub in MVB sorting at the level of cargo recognition (Bilodeau et al., 2002; Katzmann et al., 2001; Reggiori and Pelham, 2001) as well as the dependence of ILV formation on the presence of ubiquitylated-cargo (MacDonald et al., 2012a; Stringer and Piper, 2011). However, upstream effects on cargo recognition and early ESCRT membrane recruitment precluded examination of the role of Ub in ILV formation itself (Stringer and Piper, 2011). The present studies indicate that Ub-binding also contributes to ILV formation as disruption of Ub-binding through point mutation (Figure 9F) or deletion (Figure S2B) reduced Bro1^ΔBOD^ NBD-PC sorting into the vacuole lumen, consistent with Ub-enhanced V stimulation of Vps4 *in vitro*. The source of Ub *in vivo* remains to be determined, but could be Ub-cargo, ubiquitylated ESCRT machinery, or perhaps free Ub released from the Bro1-associated ubiquitin-peptidase, Doa4 (Ali et al., 2013; Kalinowska et al., 2015; Luhtala and Odorizzi, 2004). Our results implicate a role for Ub in modulating Vps4 activity via Bro1 in addition to Ub’s previously appreciated role as a cargo sorting determinant.

We suggest that Bro1 serves to “license” ILV formation, typically coordinated with cargo entry, to enable efficient MVB sorting. This licensing concept is supported by our observation that highly-expressed Bro1 promoted more ILV formation, as assessed by NBD-PC sorting, than Bro1^ΔBOD^ expressed at lower levels. A broader role for V domains in the regulation of Vps4/ESCRT-III dependent functions is implied by our observation that the V domain from another member of the Bro1 Domain family, HD-PTP, also stimulated Vps4 activity. Bro1 Domain Family members have important yet unclear roles in myriad ESCRT-mediated processes. ALIX binds viral structural proteins, such as Gag, to facilitate retroviral budding as well as other factors (e.g., Syntenin, CEP55) to facilitate ESCRT-driven exosome biogenesis and cytokinesis (Baietti et al., 2012; Carlton and Martin-Serrano, 2007; Fisher et al., 2007; Larios et al., 2020; Lee et al., 2007b). Increased ALIX activity through over-expression or activating truncation enhance the release of HIV particles as well as small extracellular vesicles (Fisher et al., 2007; Larios et al., 2020), consistent with ALIX serving as a licensing factor for these ESCRT-mediated events as well. Mutations in HD-PTP are linked to a congenital neurodevelopment disorder characterized by seizures and spasticity (Bend et al., 2020; Sowada et al., 2017). Moreover, HD-PTP homozygous mouse knockouts are embryonic lethal, and HD-PTP heterozygosity is linked to increased tumorigenesis (Gingras et al., 2009; Manteghi et al., 2016). In yeast, loss of Bro1 disrupts MVB sorting and ILV formation leading to disrupted endosomal morphology (i.e., “class E vps phenotype”). Thus, understanding the biochemical mechanism that Bro1 uses to control Vps4 has wide ranging ramifications and is likely to describe a shared function for other Bro1 Domain Family proteins (Vietri et al., 2020).

## Methods

### Plasmids and Strains

For a complete listing of plasmids utilized in this study see Supplemental Table 1. All restriction enzymes were obtained from New England BioLabs Inc. (Ipswich, MA). All PCR reaction were performed using Platinum SuperFi II PCR Master Mix (Thermo Fisher Scientific, Waltham, MA). *S. cerevisiae* Vps4 MIT domain was amplified from SEY6210 genomic DNA, and subsequently cloned into *NdeI* and *SalI* sites of pET28a to generate pET28-MIT. Similarly, MIT domain was amplified from pVPS4(I18D) and pVSP4(L64D) and subsequently cloned into pET28a to generate pET28b-MIT I18D and pET28b-MIT L64D. The sequence encoding the V domain of Bro1 (Bro1V, aa 370-709) was amplified from pGO216 (Wemmer et al., 2011) and cloned into the *BamH1* and *Sal*I sites of pGST-parallel1 (Sheffield et al., 1999) to generate pSD1. *BRO1* was amplified from pGO187 (Odorizzi et al., 2003) and cloned into the *SpeI* and *Sal*I site of pRS414 (Simons et al., 1987) to generate pSD12. Mutagenesis of Bro1V or *BRO1* encoding constructs (pSD2-9, pSD13-14, pCT2, pCT4-8, pCT10, pCT13-16, pCT18, pCT20-24, pCT26, and pCT29-33 was performed using the GeneTailor Site-Direct Mutagenesis System (Life Technologies, Carlsbad, CA). The veracity of plasmids was confirmed by sequencing. The sequence encoding Bro1ΔPRR (aa 1-709) was amplified from pGO216 and cloned into *Bam*H1 and *Sal*I sites of pET28b (Novagen, Germany) and pGST-parallel1 for use generating and purifying the anti-Bro1 sera, respectively (see *Bro1 antibody generation*).

Two silent mutations (D133D, D153D) were introduced into the *HIS3* cassette of pRS413 to remove undesired *Bgl*II sites by site-directed mutagenesis; the resulting plasmid was named pCT34. The sequence encoding *HIS3* cassette was amplified from pCT34 by PCR to introduce *Bam*HI and *Pac*I sites, and cloned into *Bgl*II and *Pac*I sites of pFA6a-HisMX6; the resulting plasmid is named pCT35. The sequence encoding Bro1V (aa370-709) was amplified from SEY6210 genomic DNA, and cloned into *Nco*I and *EcoR*I sites of pCT35; the resulting plasmid is named pCT36. Subsequently, 500bp of *BRO1* 3’UTR was amplified from SEY6210 genomic DNA and cloned into *EcoR*I and *Sac*I sites of pCT35; the resulting plasmid is named pCT37. *BRO1* promoter was amplified from genomic DNA of SEY6210, and cloned into *Sal*I and *EcoR*I sites of pRS414; the resulting plasmid is named pCT38. pCT38 generated by deleting BOD (aa1-369) in pSD12 using GeneTailor Site-Direct Mutagenesis System. Subsequently, ΔUBD mutations (I377R L386R) were introduced by site directed mutagenesis; the resulting plasmid is named pCT39. Fragment containing M8 mutation was cloned into *Cla*I and *Nde*I site of pCT38 from pSD14; the resulting plasmid is named pCT40. *TEF1*p-Bro1ΔBOD (aa370-844) was amplified from CTY2 and cloned into *Sal*I and *Bam*HI sites in pRS414; the resulting plasmid is named pCT41. Mutations (ΔUBD and M8) were amplified from pCT39 and pCT40 and cloned into *Nco*I-*Bam*HI digested pCT41 by NEBuilder HiFi DNA Assembly (New England BioLabs Inc.). DPAP-B-GFP was cloned from pGO89 (Odorizzi et al., 1998) into *Not*I and *Sal*I sites of pRS425; the resulting plasmid is DPAP-B-GFP. For a list of primers used to generate pCT34-43, please see Supplemental Table 2.

For a complete listing of strains used in this study see Supplemental Table 3. CTY1, CTY2, and CTY3 were generated by transforming PCR product from pCT34 into SEY6210.1 while CTY4 was generated by transforming PCR product from pCT36; for a list of primers used, please see Supplemental Table 2. Transformants were plated on Histidine drop-out synthetic plates, and screened by immunoblotting. The altered *bro1* allele was amplified and sequenced. CTY5, CTY11-13, CTY518, CTY21-22, CTY24, CTY27, CTY29-30 and JPY403 were generated using standard yeast genetics.

### Protein expression and purification

Protein expression was performed in the BL21-DE3 bacterial strain using 0.5 mM isopropyl ß-d-1-thiogalactopyranoside (IPTG) at 16°C for 18-22hrs. GST-Bro1V fusion protein and mutants thereof were purified using the Glutathione Sepharose 4B (GE Healthcare, United Kingdom), treated with AcTEV protease (Thermo Fisher Scientific, Waltham, MA) overnight at room temperature to remove the GST tag, incubated with ATP to dissociate chaperones, and subjected to size-exclusion chromatography (Superdex200 HiLoad 16/60) in 25mM HEPES and 150mM KCl. Purified proteins were tested for purity using SDS-Page and Coomassie staining and were analyzed to exclude contaminating ATPase activity (data not shown). All Bro1V mutants examined in this study exhibited equivalent expression levels in bacteria (data not shown) and mutants expressed in yeast were at levels similar to WT (Figure 6b). GST-Vps4 was expressed and purified as previously described for use in the ATPase assays (Babst et al., 1997). His_6_-Vps4 was purified as previously described (Davies et al., 2014), except the His_6_ fusion tag was not cleaved in order to use His_6_-Vps4 in the *In vitro* Binding Assays. His-Bro1ΔAPRR purified from BL21-DE3 using Ni^2+^-affinity chromatography (5 ml HiTrap Chelating FF column; GE Healthcare, United Kingdom). GST-Bro1V(370-709) was purified using the Glutathione Sepharose 4B (GE Healthcare, United Kingdom) and eluted with reduced glutathione (Thermo Fisher Scientific, Waltham, MA). V domains of *S. castellii* Bro1 and human HD-PTP were produced in *E. coli* BL21 (DE3) were purified by TALON-Co^2+^ and size exclusion chromatography and cleaved from their 6xHis tag with AcTEV protease (Pashkova et al., 2013). Mono-Ub was expressed in *E. coli* BL21 (DE3) from pPL5293. The protein was purified from bacterial cell lysate by precipitation with 5% Perchloric acid and CMC cation exchange chromatography in 0.05M ammonium acetate buffer pH 4.5 followed by elution with 0.1M ammonium acetate buffer pH 5.5 (Sundd et al., 2002).

### Antihody generation

Anti-Snf7 antiserum was generated against GST-Snf7. Antiserum was generated in a New Zealand rabbit (Covance, Princeton, NJ). Bleeds were evaluated for detection of Snf7 (SEY6210 and *snf7Δ*). The Snf7 antibody was used at 1:5,000. Anti-Bro1 antiserum was generated against His_6_-Bro1ΔPRR. Antiserum was generated in a New Zealand rabbit (Covance, Princeton, NJ). Antibodies were purified using GST-Bro1ΔPRR immobilized on an AminoLink column (Thermo Fisher Scientific, Waltham, MA) and verified for sensitivity and specificity using yeast extracts (SEY6210 and *bro1*Δ) and recombinant Bro1V. The purified Bro1 antisera was used at 1:1,000.

### *In vitro* binding assays

Ni-NTA Agarose beads (Qiagen, Germany) +/- His_6_^-^Vps4 (10μg) was incubated for 1 hr at 4°C in NiA buffer (25mM NaH_2_PO_4_, 300mM NaCl, pH 7.5), washed with NiA buffer, and equilibrated with ATPase buffer (see *ATPase assay*). Binding reactions were performed in Handee Spin Columns (Pierce) with 6μg Bro1V or mutants in ATPase buffer + 0.02% Tween20 incubated for 1 hour at 30°C. Reactions were washed four times with NiA buffer with 10mM imidazole (Sigma Aldrich, St. Louis, MO), and the bound material was eluted with NiA buffer with 200mM imidazole. Instant-Bands Protein Stain (EZBiolab, Parsippany, NJ) were used to visualize input and eluted material on a Typhoon FLA 7000 (GE Healthcare, United Kingdom). Samples were resolved via SDS-PAGE, and detected using immunoblotting against Bro1. Bro1V input and His_6_-Vps4 were detected via SDS-PAGE followed by Coomassie staining (data not shown).

Glutathione Sepharose 4B GST-tagged protein purification resin (Cytiva Life Sciences, Marlborough, MA) +/- lysate from *E. coli* BL21 (DE3) expressing GST-tagged Bro1 fragments were incubated for 1hr at 4°C in PBS, washed with PBS and equilibrated with ATPase buffer. Binding reactions were performed in Handee Spin Columns with His_6_^-^Vps4 in ATPase buffer + 0.02% Tween20 incubated for 1 hour at 30°C. Reactions were washed six times with PBS + 0.02% Tween20, and the bound material was eluted with GST elution buffer (10mM reduced glutathione, 50mM Tris-HCl, pH8.0). Instant-Bands Protein Stain were used to visualize input and eluted material on a Typhoon FLA 7000. Samples were resolved via SDS-PAGE, and His_6_^-^ Vps4 was detected using Penta-His Antibody (Qiagen, Germany) per manufacturer instruction. Three or more independent experiments were performed with each experiment performed in replicates within the experiment and representatives are shown.

### ATPase assays

Measurement of Vps4 ATPase activity was performed in ATPase buffer (20 mM HEPES, 100 mM KOAc, 5mM MgOAc, pH 7.5) as previously described (Azmi et al., 2008; Babst et al., 1998; Davies et al., 2010; Davies et al., 2014; Norgan et al., 2013; Tan et al., 2015). All reactions were incubated at 30°C for 30 minutes prior to initiation by ATP addition (4mM final concentration). Images were captured using a Typhoon FLA 7000 (GE Healthcare, United Kingdom). Vps4 titration data represent ATPase activities from a minimum of three independent experiments with each experiment performed in duplicates. Data represent ATPase activities from a minimum of three independent experiments with each experiment performed in replicates within the experiment. Data was graphed and statistical significance was assessed by T tests using Prism 5 (GraphPad, San Diego, CA). The Vps4 concentration used was 0.5μM because this concentration of Vps4 exhibits sub-maximal specific activity. ATPase assays in the presence of ubiquitin were performed with the minimal concentration (50μM) yielding maximal enhancement.

### ESCRT-III disassembly assays

ESCRT-III disassembly assays were performed as previously described (Davies et al., 2010; Tan et al., 2015). Briefly, spheroplasted yeast (*vps4*Δ *pep4*Δ *bro1*Δ) were lysed with a Dounce homogenizer in 20mM HEPES, 200mM Sorbitol, 50mM KOAc, 2mM EDTA, pH 6.8, and subjected to a 5 minute, 500xg clearing spin. The resulting supernatant was then spun 10 minutes at 13,000xg. Using Disassembly Assay buffer (20mM HEPES, 100mM KOAc, 5mM MgOAc, 200mM Sorbitol, pH 7.4 with protease inhibitors), the pelleted membranes were washed and then resuspended at 50 OD_600_ equivalents/ml. Reactions were setup with 0.5 OD_600_ equivalents/ml, an ATP regeneration system (1mM ATP) and various amounts of Vps4 and Bro1V. Vps4 and Bro1V were allowed to preincubate prior to reaction initiation. The reactions were initiated by brief centrifugation followed by incubation at 30°C for 10 min. Reactions were terminated by centrifugation at 13,000xg for 10 min. immunoblotting analysis was used to assess the levels of Snf7 in the pellet and soluble fractions. Snf7 was detected using anti-Snf7 polyclonal antibody (1:5000), HRP-conjugated Goat anti-Rabbit IgG (Life Technologies, Carlsbad, CA, 1:30,000) and the UVP Autochemi System (Upland, CA) and quantified using ImageQuant software (GE Healthcare, United Kingdom). Linearity of the signal was confirmed via image histograms. Disassembly assay data represent three independent experiments with each experiment performed in replicates within the experiment. Data was graphed and statistical significance was assessed by T tests using Prism 5 (GraphPad, San Diego, CA). Data are graphed as the mean with the standard error of the mean, and representative immunoblots are shown.

### Fluorescence microscopy

For live-cell imaging of cells expressing GFP-tagged cargoes were grown in minimal media at 30°C to mid-log phase (0.5OD_600_) for live-cell fluorescence microscopy. Mup1-GFP induction was accomplished by adding 50μM copper sulfate/chloride, as previously described (MacDonald et al., 2015). Co-labelling with FM4-64 and C6-NBD-PC (1-palmitoyl-2-(6-((7-nitro-2-1,3-benzoxadiazol-4-yl)amino)hexanoyl)-sn-glycero-3-phosphocholine; Avanti Polar Lipids, Alabaster, AL) was performed by growing cells in YPD or Synthetic Media containing 1μM C6-NBD-PC and 250nM FM4-64 for 30 minutes at 30°C. Cells were then harvested at mid-log phase and imaged. Images were captured at room temperature using an Olympus IX70-S1F2 fluorescence microscope (Center Valley, PA) equipped with an Olympus UPIanApo 100x numerical aperture 1.35 oil objective with the complementing immersion oil (N=1.516, Applied Precision, Issaquah, WA), Standard DeltaVision filters FITC and Rhodamine, and Photometrics CoolSNAP HQ CCD monochrome camera (Teledyne photometrics, Tucson, AZ). Image was acquired using Delta Vision softWoRx (version 3.5.1, Applied Precision, Issaquah, WA) and subsequently processed by FiJi (version: 2.1.0/1.53c, National Institutes of Health, Bethesda, MD) (Schindelin et al., 2012). Captured images were exported under the standard DeltaVision file format and converted into 16-bit TIFF images by using Bio-Formats Importer available within Fiji. The contrast and brightness of images were subsequently adjusted within Fiji as well. Images were captured on three different days using two different sets of transformations. Verification of Bro1 and mutant Bro1 expression in these cells was confirmed by immunoblotting. Cells with WT cargo sorting signal were scored manually. Data represent quantification from a minimum of three independent labeling experiments with each experiment quantified at least 100 cells. Statistical significance was assessed by T tests using Prism 5 (GraphPad, San Diego, CA).

### Cargo GFP liberation analysis

Cells expressing GFP-tagged cargoes grown in minimal media at 30°C to mid-log phase (0.5OD_600_) were harvested and treated with 0.2M sodium hydroxide for 5 minutes. Subsequently, cells were resuspended in urea buffer (40 mM Tris-HCl, pH 6.8, 8 M urea, 15% SDS w/v, 0.1 mM EDTA, 1% β-mercaptoethanol v/v, 0.01% bromophenol blue w/v) and lysed with glass bead for 5 minutes (Katzmann et al., 1999). Sample were resolved via SDS-PAGE and immunoblotted for GFP (Roche AG, Basel, Switzerland, 1:1,000) and Pgk1 (Life Technologies, Carlsbad, CA, 1:10,000).

### Subcellular fractionations

Subcellular fractionations were performed by osmotic lysis as previously described (Dimaano et al., 2007), except the cell extracts were centrifuged at 13,000xg for 10 min at 4°C to separate the soluble and pellet fractions (Tan et al., 2015). Samples (0.1 or 0.2 OD_600_ equivalents) were resolved via SDS-PAGE and immunoblotted for Snf7 (this study, 1:5,000), Vps4 ((Babst et al., 1997), 1:1000), Bro1 (this study, 1:1,000). Pgk1 (Life Technologies, Carlsbad, CA, 1:10,000), and Pep12 (Life Technologies, Carlsbad, CA, 1:1,000). Immunoblots were developed using HRP-conjugated Goat anti-Rabbit (Life Technologies, Carlsbad, CA, 1:30,000) or Goat anti-Mouse (Life Technologies, Carlsbad, CA, 1:1,000), SuperSignal West Pico and SuperSignal West Femto substrates (Thermo Fisher Scientific, Waltham, MA), and the UVP Autochemi System (Upland, CA). Signal was quantified using ImageQuant software (GE Healthcare, United Kingdom). Signal linearity was confirmed via image histograms. Data represent three or more independent experiments (representative immunoblots are shown) and are graphed as mean with standard error of the mean. Statistical significance was assessed by T tests using Prism 5 (GraphPad, San Diego, CA).

### Statistical Analysis

Statistical significance was assessed by parametric unpaired two-tailed Student’s T test.

### Molecular modeling

Molecular graphics and analyses of the *S. castellii* Bro1 V domain (Protein Data Bank ID: 4JIO, chain A) were performed with the UCSF Chimera package and PyMOL (Pettersen et al., 2004; Schrödinger). Chimera was developed by the Resource for Biocomputing, Visualization, and Informatics at the University of California, San Francisco (supported by NIGMS P41-GM103311).

### Dual axis electron tomography

Yeast cells were high-pressure frozen (HPF) and freeze-substituted (FS) as previously described (Buysse et al., 2020; Johnson et al., 2017). Liquid cultures were harvested during log phase and filtered with 4.5-micron Millipore paper, collected into 0.5 mm aluminum hats, high pressure frozen with a Wohlwend HPF (Wohlwend, Switzerland), and transferred to FS media kept at liquid nitrogen temperature until cryo-fixation. Cells were then freeze-substituted in an AFS (Automated Freeze-Substitution machine, Leica Vienna, Austria) at −90°C in cryo media made from 0.1% uranyl acetate 0.25% glutaraldehyde in anhydrous acetone, washed in pure acetone, and embedded at −60°C in Lowicryl HM20 (Polysciences, Warrington, PA). Samples were polymerized at −60°C with and warmed slowly over four days. Plastic blocks were trimmed, and sections were cut in 80 nm thin sections and 250 nm thick sections with a Leica UC6 ultramicrotome and placed on Rhodium-plated copper slot grids (Electron Microscopy Sciences, Hatfield, PA) for thin section transmission electron microscopy (TEM) and thick section electron tomography (ET). TEM of hundreds of cells per strain is used to quality control freezing, embedding, and staining for tomography (Richter et al., 2013). Tomographic samples are en bloc stained with 0.1% uranyl acetate and 0.25% glutaraldehyde only, with no additional post staining as before (Giddings, 2003). 100 cells were surveyed before at least 10 representative cells were selected to gather the tomography.

Thick sections were labeled with 15 nm fiducial gold on both sides and mapped on a Phillips CM10 (TEM; Mahwah, NJ) at 80-kv and tilt-imaged with a Tecnai 30 (300kV FEI; Beaverton, OR) with dual tilt series images collected from +60 to −60 degrees with 1 degree increment using a Gatan US4000 4k x 4k CCD camera. Tilt-series were shot at 19,000 times magnification with a 0.6534 nm working pixel (binning 2) and repeated at a 90 degree rotation for dual-axis tomography. Tomograms and models were constructed using IMOD software package (Kremer et al., 1996).

MVB membrane models from dual-axis electron tomograms are manually assigned from the inner leaflet every 4 nm and calculated using Imodmesh (3DMOD). We designated budding profiles (BPs) by their negative curvature since the majority of endosome limiting membrane curvature is positive, or spherical in shape. BP models are drawn from the 0° rim at the outer leaflet, measured, and sorted by surface area using only BPs that have more than 750 nm^2^ or approximately half of the mean ILV surface (Wemmer et al., 2011). ILVs are spherical and measured using sphere-fitting models from the vesicle’s outer leaflet (the inner leaflet of the MVB limiting membrane) and ILV diameters are measured using these sphere models. Movies were made using IMOD and QuickTime (Apple, Cupertino, CA). Data were analyzed and graphed using Prism 5 (GraphPad, San Diego, CA).

## Acknowledgements

We would like to acknowledge Drs. Gina Razidlo and Jason Tan (Mayo Clinic, Rochester) for their inquiries and thoughtful discussions. SD, CT contributed to these studies as members of the Biochemistry and Molecular Biology Graduate Program and received financial support from the Mayo Clinic Graduate School of Biomedical Sciences. JS contributed to these studies as a member of the Mayo Clinic Summer Undergraduate Research Program and received financial support from the Mayo Clinic Graduate School of Biomedical Sciences. The authors declare no competing financial interests.

## Author Contributions

Brian A. Davies, Ishara F. Azmi and David J. Katzmann conceived of the study. Chun-che Tseng, Shirley Dean, Brian A. Davies, and David J. Katzmann designed and conducted the experiments along with Johanna A. Payne, Jennifer Staffenhagen, Natalya Pashkova, Robert C. Piper, Matt West, and Greg Odorizzi. David J Katzmann, Robert C. Piper, Chun-che Tseng, and Brian A. Davies wrote the manuscript. All authors reviewed the results and approved the final version of the manuscript.

---

**Figure S1. Representative images of cells stained with NBD-PC and FM4-64.** This figure complements Figure 3. **(A)** Three-dimensional models reconstructed from 200-nm-thick section electron tomograms of *bro1*Δ (GOY65) cells. This image depicts rare MVBs that contains an ILV. Normal like endosomes are highlighted by yellow limiting membrane, while other colors depict flattened or tubular endosomes devoid of vesicles. The ILVs are highlighted in red. Scale bar = 100nm. **(B)** Selection criteria for scoring positive NBD-PC cells. A usable cell is defined by having (1) readily identifiable FM4-64 labeling of vacuole membrane, (2) readily identifiable NBD-PC signal, and (3) defined vacuole(s). An unusable cell is defined by having (1) a fragmented vacuole, and (2) out of focus. A positive cell is defined by a diffuse NBD-PC signal within the lumen while lacking a distinct ring on the limiting membrane of the vacuole (defined by FM4-64); while a negative cell is defined by the colocalization of NBD-PC and FM4-64 on the limiting membrane of the vacuole. Scale = 5μm. **(C)** Representative micrographs of *bro1*^Δ*BOD*^ (*bro1*Δ::*TEF1*p-*bro1*^Δ*BOD*^ CTY2) and *bro1*^Δ*BOD*^ *vps4*Δ (CTY5) cells stained with NBD-PC to label ILVs and FM4-64 to label vacuoles. Scale = 5μm.

**Figure S2. *bro1*^Δ*BOD*^ supported ILV formation requires the Vps4/ESCRT machinery.** This figure complements Figure 3. **(A)** Lysates generated from WT (SEY6210), *vps4Δ* (MBY3), *bro1*Δ (GOY65), *TEF1p-BRO1* (CTY1), *bro1*^Δ*BOD*^ (370-844; *bro1*Δ::*TEF1*p-*bro1*^Δ*BOD*^; CTY2), *bro1*^Δ*BOD*^ (Δ*UBD*) (388-844; *bro1*Δ::*TEF1*p-*bro1*^Δ*BOD*^ CTY3), *TEF1*p-*bro1V*(CTY4), and *bro1*^Δ*BOD*^ *vps4*Δ (CTY5) cells were analyzed by immunoblotting using antibodies against Bro1, Vps4 and Pgk1. Numbers below the Bro1 blot indicates expression levels normalized to *BRO1* expression. **(B)** WT (SEY6210), *bro1*^Δ*BOD*^ (370-844; CTY2), *bro1*^Δ*BOD*,Δ*UBD*^ (388-844; CTY3), *bro1*^Δ*BOD*^ *hse1*Δ (CTY30), *bro1*^Δ*BOD*^ *vps27*Δ (CTY29), *bro1*^Δ*BOD*^ *vps37*Δ (CTY21), *bro1*^Δ*BOD*^ *mvh12*Δ (CTY22), *bro1*^Δ*BOD*^ *vps22*Δ (CTY24), *bro1*^Δ*BOD*^ *snf7*Δ (CTY12), *bro1*^Δ*BOD*^ *vps24*Δ (CTY18), *bro1*^Δ*BOD*^ *vps2*Δ (CTY26), *bro1*^Δ*BOD*^ *vta1*Δ (CTY27), *bro1*^Δ*BOD*^ *vps4*Δ (CTY5), and *bro1*^Δ*BOD*^ *doa4*Δ (CTY13) cells were analyzed by live cell fluorescence microscopy and quantified for the frequency of cells able to support NBD-PC trafficking to the vacuolar lumen. Error bars indicate S.E.M. **(C)** WT (SEY6210), *bro1V* (CTY4) were analyzed by electron tomography and quantified to assess ILV size (diameter). A minimum of 22 MVBs and 225 ILVs from at least 10 cells were quantified. Error bars indicate S.E.M. * indicates statistically significant differences compared to WT. **(D)** WT (SEY6210), *bro1*Δ (GOY65), or *bro1V (bro1Δ::TEF1p-bro1V;* CTY4) cells were transformed with the indicated GFP-tagged cargo plasmid to assess MVB sorting using live cell fluorescence microscopy. White dashed lines indicate cell boundaries. Scale bar = 5μm. **(E)** WT (SEY6210), GOY65, and GOY65 cells transformed with *BRO1* and *BRO1*p-*bro1*^Δ*BOD*^ was analyzed by live cell fluorescence microscopy and quantified for the frequency of cells able to support NBD-PC trafficking to the vacuolar lumen. Error bars indicate S.E.M. **(F)** Lysates generated from *bro1*Δ (GOY65) transformed with empty vector, *BRO1*, and *BRO1*p-*bro1*^Δ*BOD*^, were analyzed by immunoblotting using antibodies against Bro1 and Pgk1.

**Figure S3. Bro1V mutants bind Vps4.** This figure complements figure 5 and 6. **(A)** Immobilized His_6_-Vps4 or Ni-NTA beads alone were incubated with Bro1V or Bro1V mutants and bound material was analyzed by immunoblotting with anti-Bro1 antiserum. **(B)** Mutant protein expression levels normalized to WT. Bro1 expression levels were normalized to Pgk1, and subsequently normalized to WT/Pgk1 ratios. Quantified from three independent experiments performed on three different days from two sets of transformations. Immunoblots were probed against Bro1 and PGK1, using lysates of GOY65 transformed with empty plasmid (pRS414) or *BRO1, bro1*(M4), *bro1*(M8), *bro1*(M9), or *bro1* (M10) plasmids. Statistical analyses did not reveal differences between WT and mutant forms of Bro1.

**Figure S4. Bro1V mutants bind ubiquitin.** This figure complements Figure 9. **(A)** Sequence alignment of V domain amino acids 370-388 from *S. castellii* and *S. cerevisiae*. Conserved amino acids are indicated by black circles, and isoleucine 377 and leucine 386 critical for Ub-binding are highlighted in red. **(B)** Bro1 V domain mutations M4, M8 and M10 (red) that disrupt V domain stimulation of Vps4 ATPase activity are spatially separated from its Ub-binding site using *S. castellii* Bro1V crystal structure (Protein Data Bank ID: 4JIO, chain A). **(C)** Vps4 titrations were performed with or without 4μM Bro1V^ΔUBD^ (L386R). Vps4 specific activity is presented. The vertical dotted line indicates the Vps4 apparent K_m_ +/- Bro1V. **(D)** Immobilized GST fused Bro1V, Bro1V(M4), Bro1V(M8), Bro1V(M9), Bro1V(M10), Bro1V(I377R) and GST alone were incubated with V5 epitope-tagged linear penta-Ub. Bound material was visualized by both Ponceau S protein stain and immunoblotting for the V5 epitope. **(E)** Lysates generated from *bro1*Δ (GOY65) transformed with empty vector, *BRO1, TEF1*p-*bro1*^Δ*BOD*^, *TEF1*p-*bro1*^Δ*BOD*,Δ*UBD*^ (Δ*UBD:I377R,L386R*) and *TEF1*p-*bro1*^Δ*BOD,M8*^ plasmids were analyzed by immunoblotting using antibodies against Bro1 and Pgk1. Numbers below the Bro1 blot indicate expression levels normalized to *BRO1* expression.

**Movie S1. Tomogram of MVB in a *bro1*Δ::TEF1p-*bro1*^Δ*BOD*^ cell.** MVBs in a *bro1*Δ::*TEFL1*p-*bro1*^Δ*BOD*^ cell exhibits WT-like MVB morphology. This video shows nine MVBs that are adjacent to the vacuole (red mesh), while the limiting membrane of the MVB is colored yellow and the intralumenal vesicles are in red. Intermediate budding profiles are highlighted in green. Scale bar = 100nm.

**Movie S2. Tomogram of MVB in a *bro1*Δ cell.** *bro1*Δ cell has class E compartments, which are flattened stacks of endosomal membranes that generally lack internal vesicles. Cisternal class E compartment stacks are shown in different colors to differentiate individual membranes. The vacuolar limiting membrane is labeled red. Scale bar = 100nm.

**Movie S3. Tomogram of MVB in a WT cell.** MVBs in WT yeast are roughly spherical membrane structures that contain smaller membrane-bound vesicles. This video shows two WT MVBs that are adjacent to the vacuole (red mesh). The limiting endosomal membrane is shown in yellow, and the intralumenal vesicles are in red. Scale bar = 100nm.

**Movie S4. Tomogram of MVB in a *bro1*Δ cell expressing Bro1^M8^.** This tomogram depicts one MVBs in a *bro1*Δ cell expressing Bro1^M8^. This MVB contains several budding intermediates (green), while its limiting membrane is indicated by yellow and the intralumenal vesicles are labeled red. Scale bar = 100nm.

**Movie S5. Tomogram of MVB in a *bro1*Δ::*TEF1*p-bro1V cell.** This tomogram depicts 14 MVBs in a *bro1Δ::TEF1p-bro1V* cell. Majority of these MVBs exhibit normal-like MVB morphology (yellow), with relatively large budding profiles (green). One tubular MVB is visible (cyan). The intralumenal vesicles are in red. Scale bar = 100nm.

**Supplemental Table 1.**
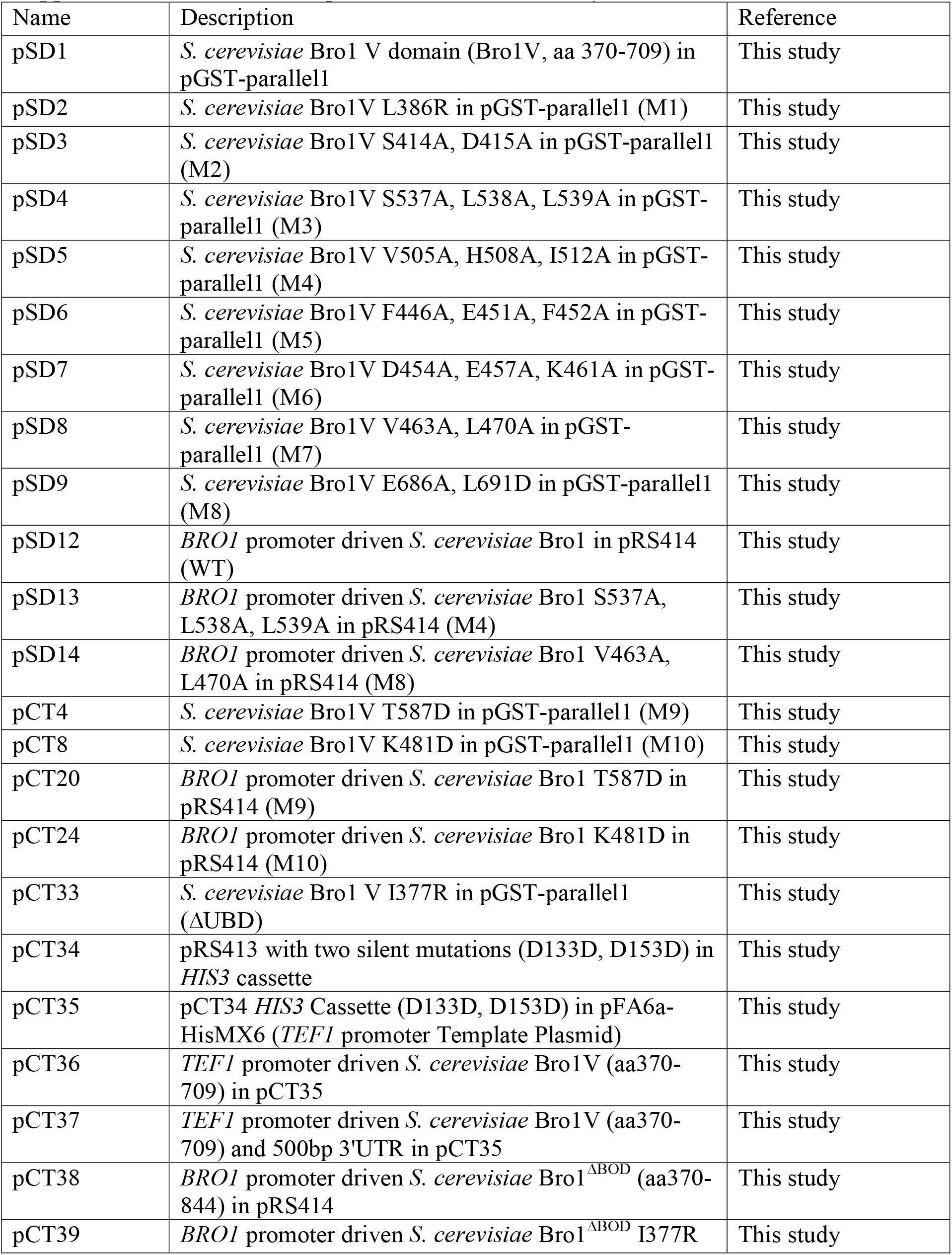
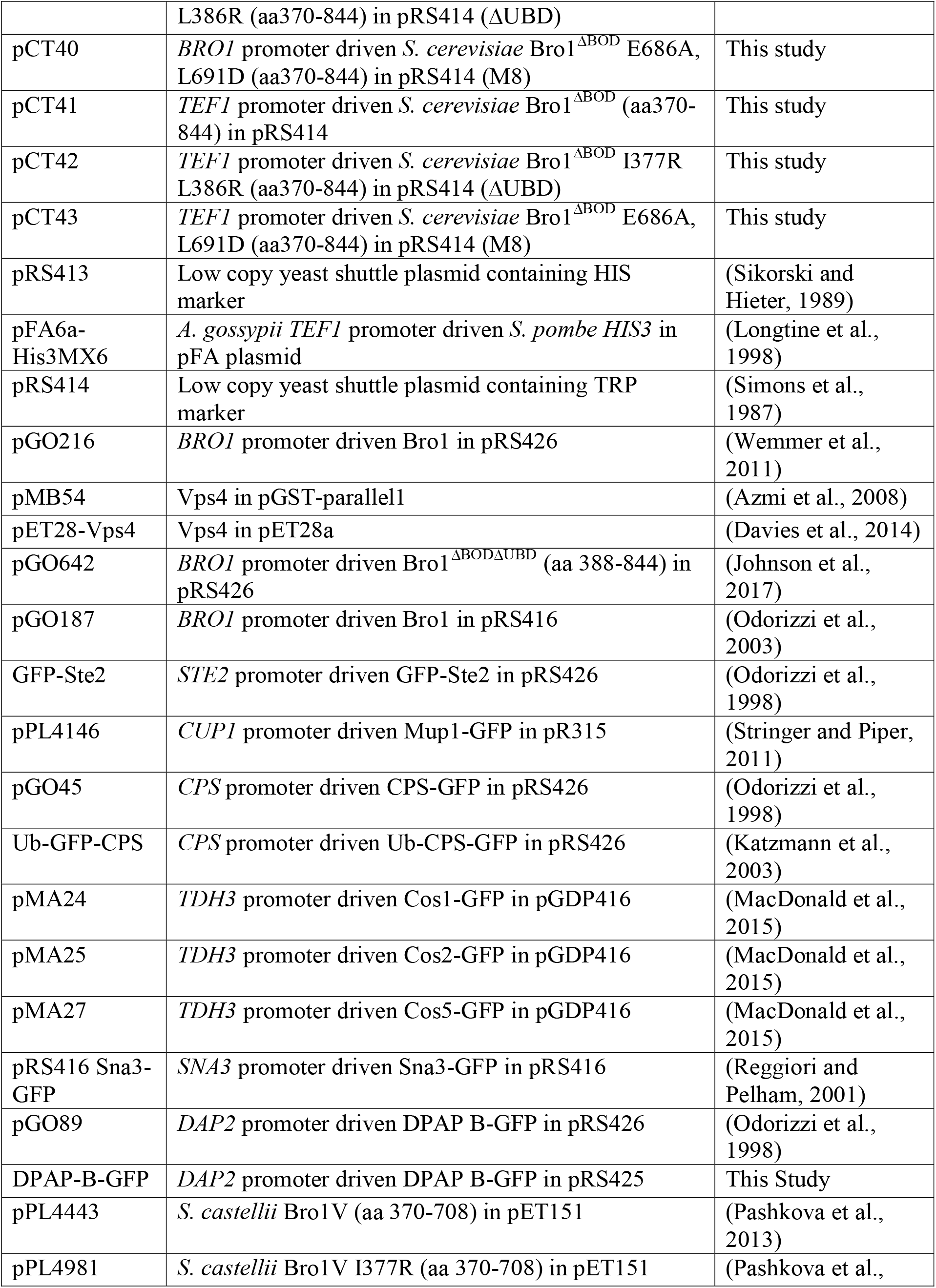
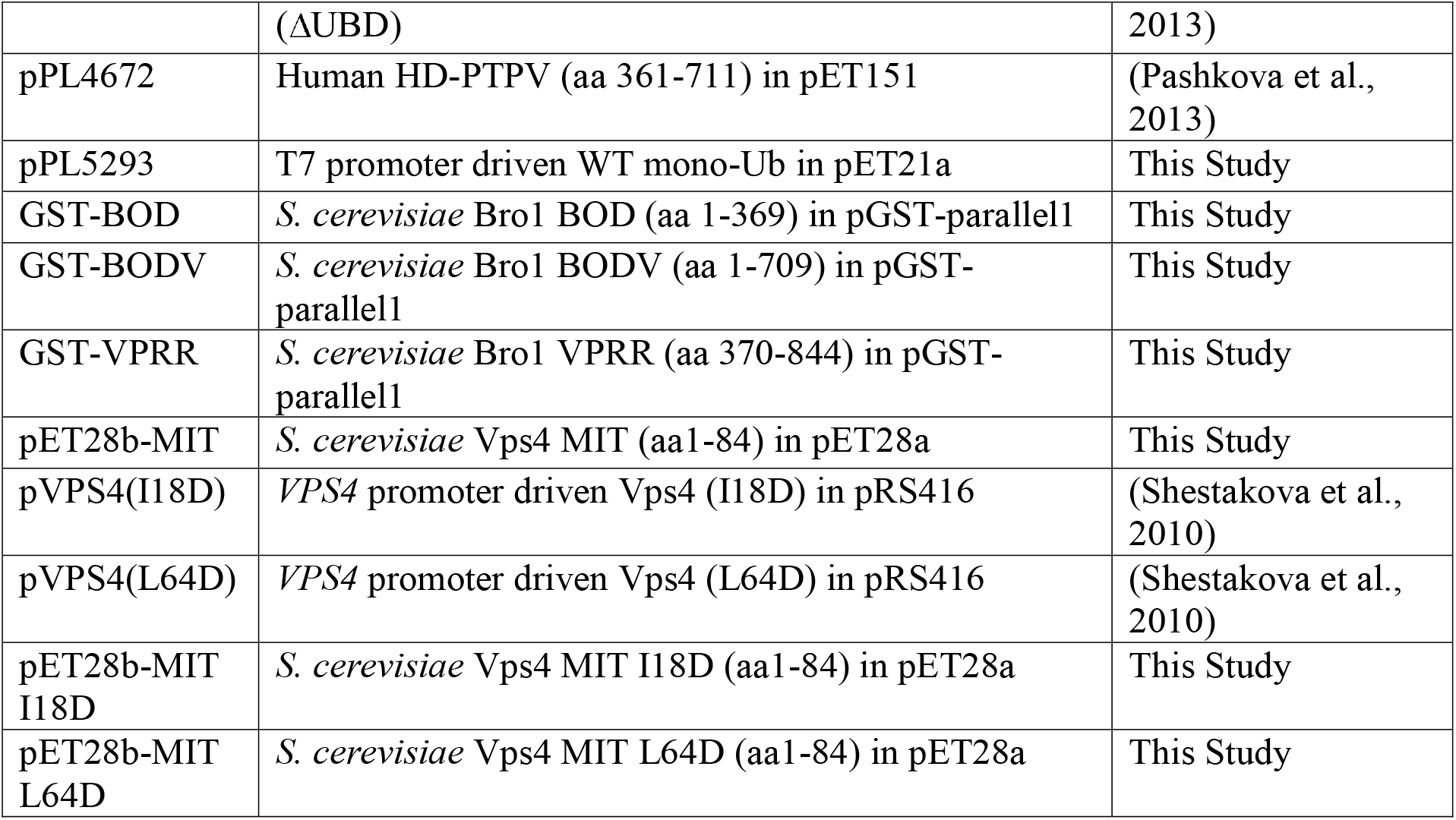
List of plasmids used in this study.

**Supplemental Table 2.**
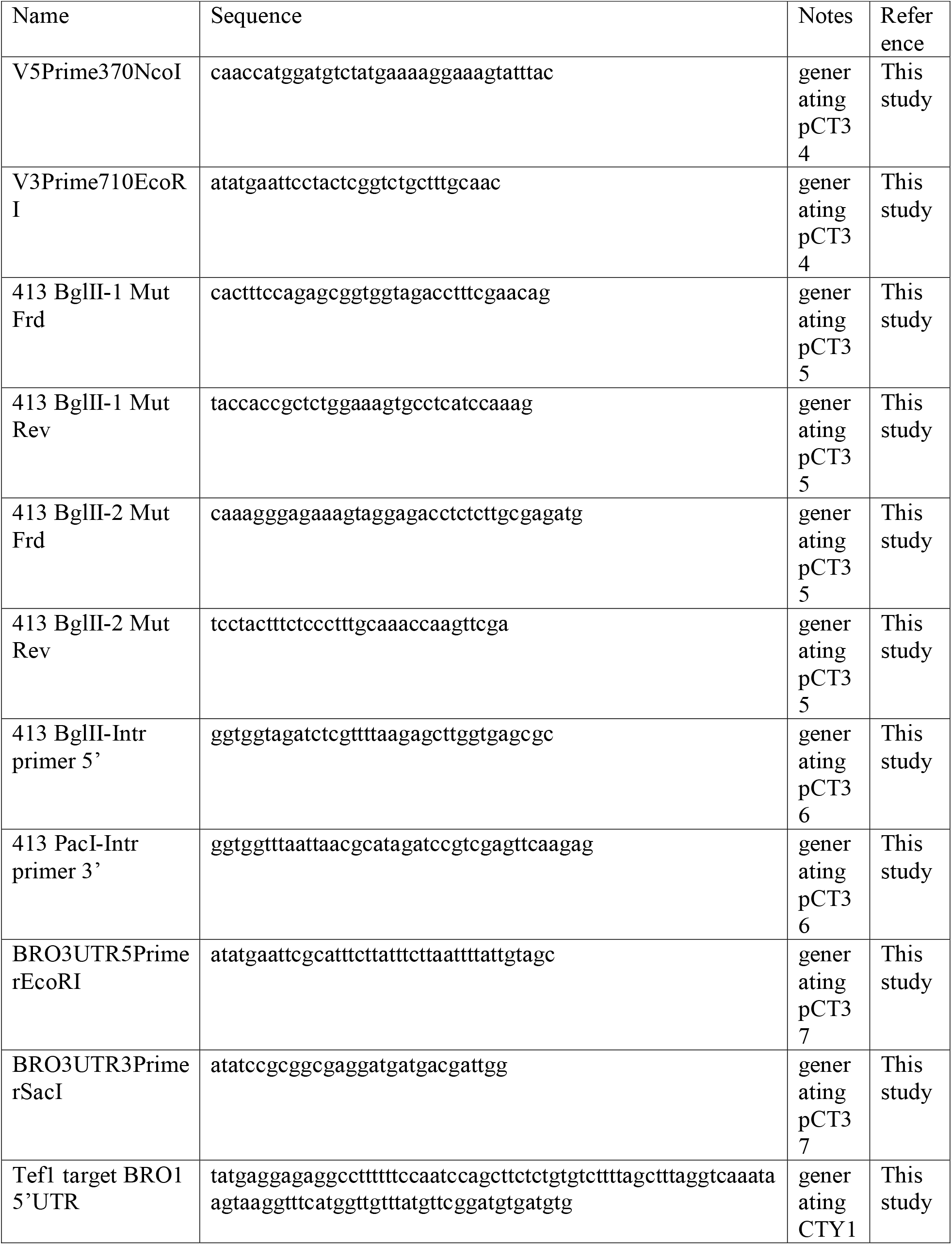
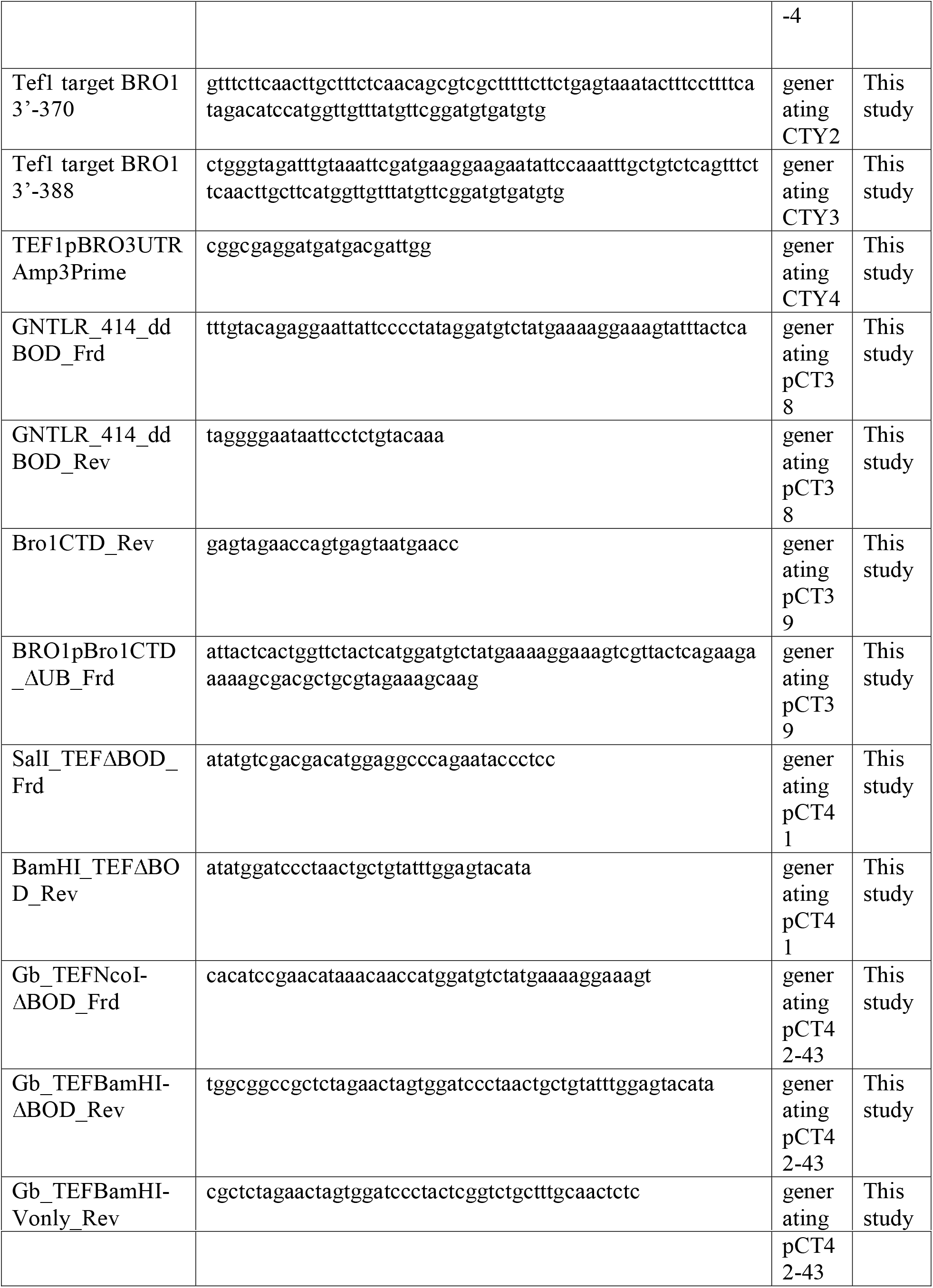
List of primers used in this study.

**Supplemental Table 3.**
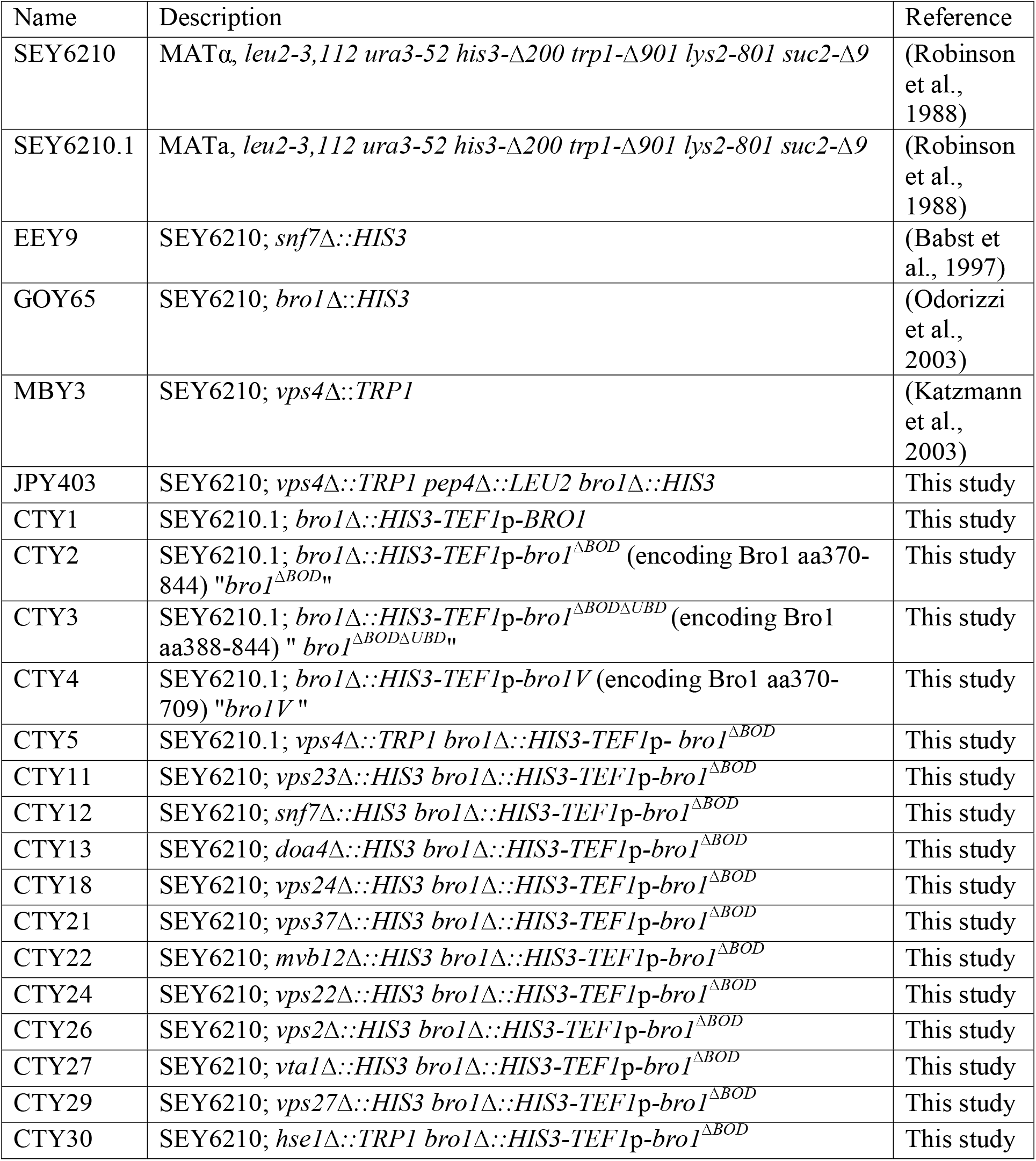
List of yeast strains used in this study.

**Supplemental Table 4.** List of mutations analyzed in this study. Only mutations that were expressed and exhibited defective ATPase stimulation or MVB cargo sorting were pursued.

| **ID** | **Mutation** | **ATPase Stimulation** | **Yeast Expression** |
| --- | --- | --- | --- |
| M1 | L386R | + | + |
| M2 | S414A, D415A | + | + |
| M3 | S537A, L538A, L539A | + | + |
| M4 | V505A, H508A, I512A | - | + |
| M5 | F446A, E451A, F452A | + | + |
| M6 | D454A, E457A, K461A | + | + |
| M7 | V463A, L470A | + | + |
| M8 | E686A, L691D | - | + |
| M9 | T587D | - | + |
| M10 | K481D | - | + |

**Abbreviations:**
ESCRT, Endosomal Sorting Complex Required for Transport; MVB, Multivesicular Body; ILV, intraluminal vesicle.

## Abbreviations

ESCRT: Endosomal Sorting Complex Required for Transport
MVB: Multivesicular Body
ILV: intralumenal vesicle.

