## Supplementary figures and images for "Bro1 stimulates Vps4 activity to promote Intralumenal Vesicle Formation during Multivesicular Body biogenesis"

### Supplemental Figure 1

#### Supplemental Figure 1

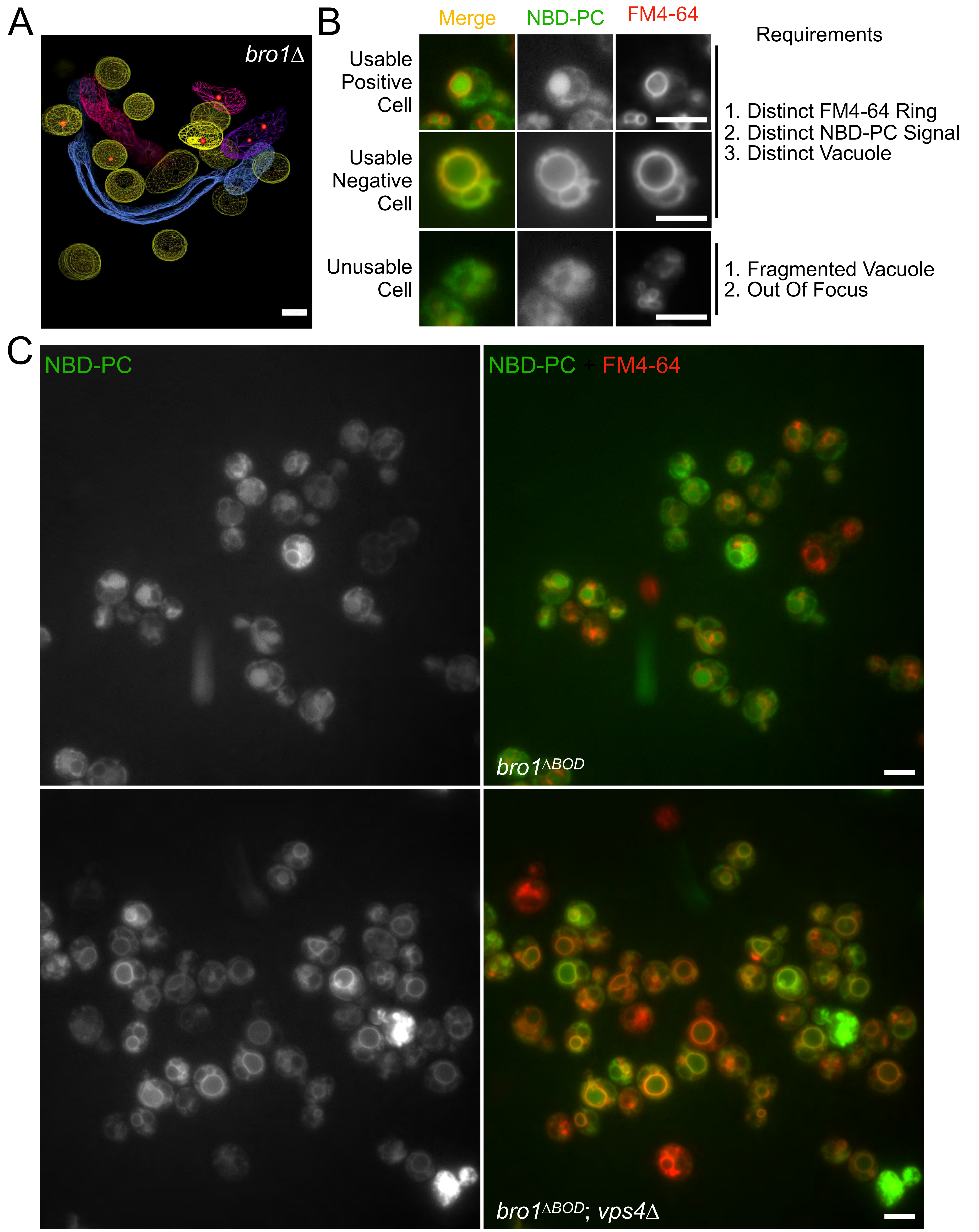

### Supplemental Figure 2

#### Supplemental Figure 2

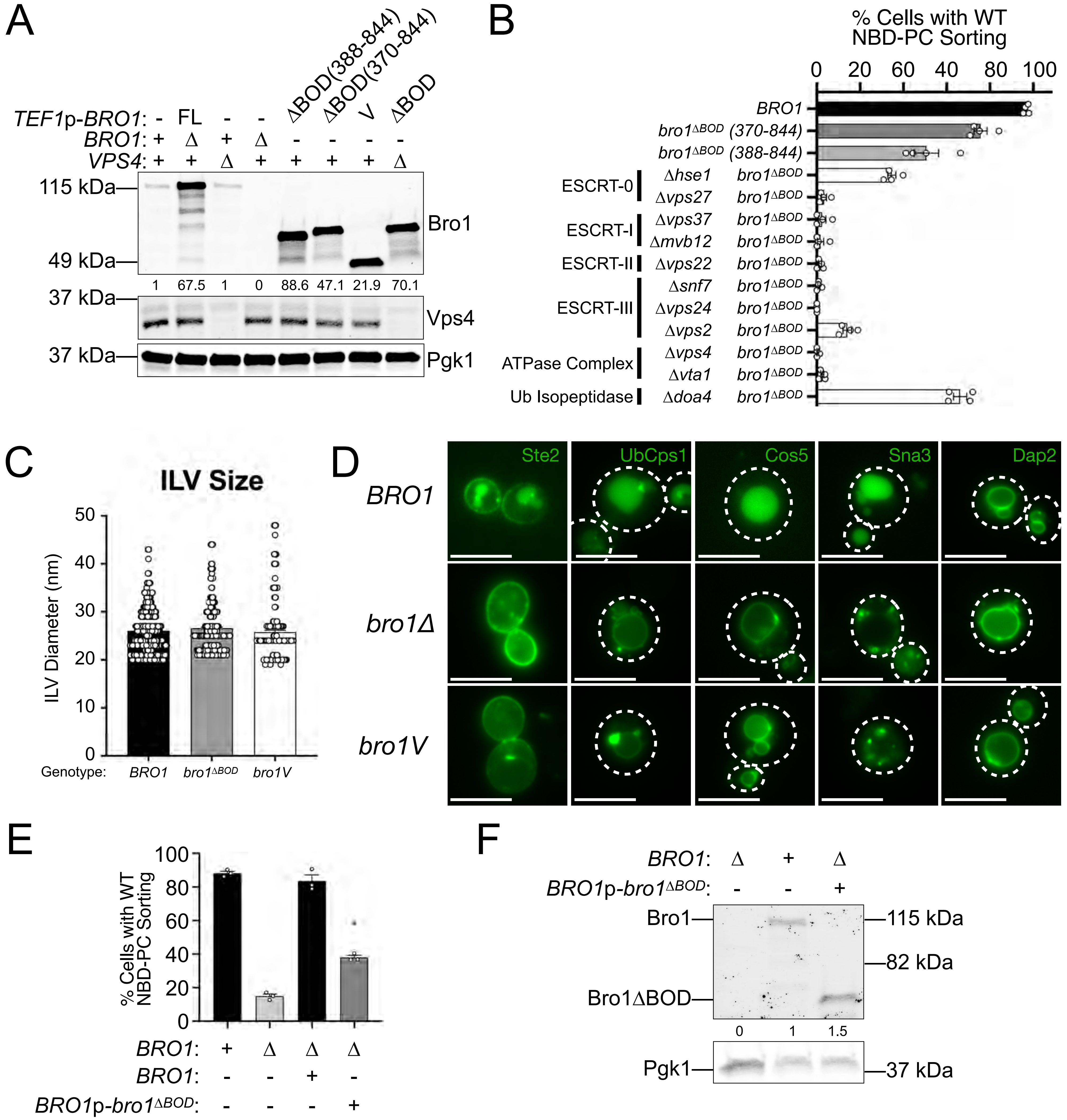

### Supplemental Figure 3

#### Supplemental Figure 3

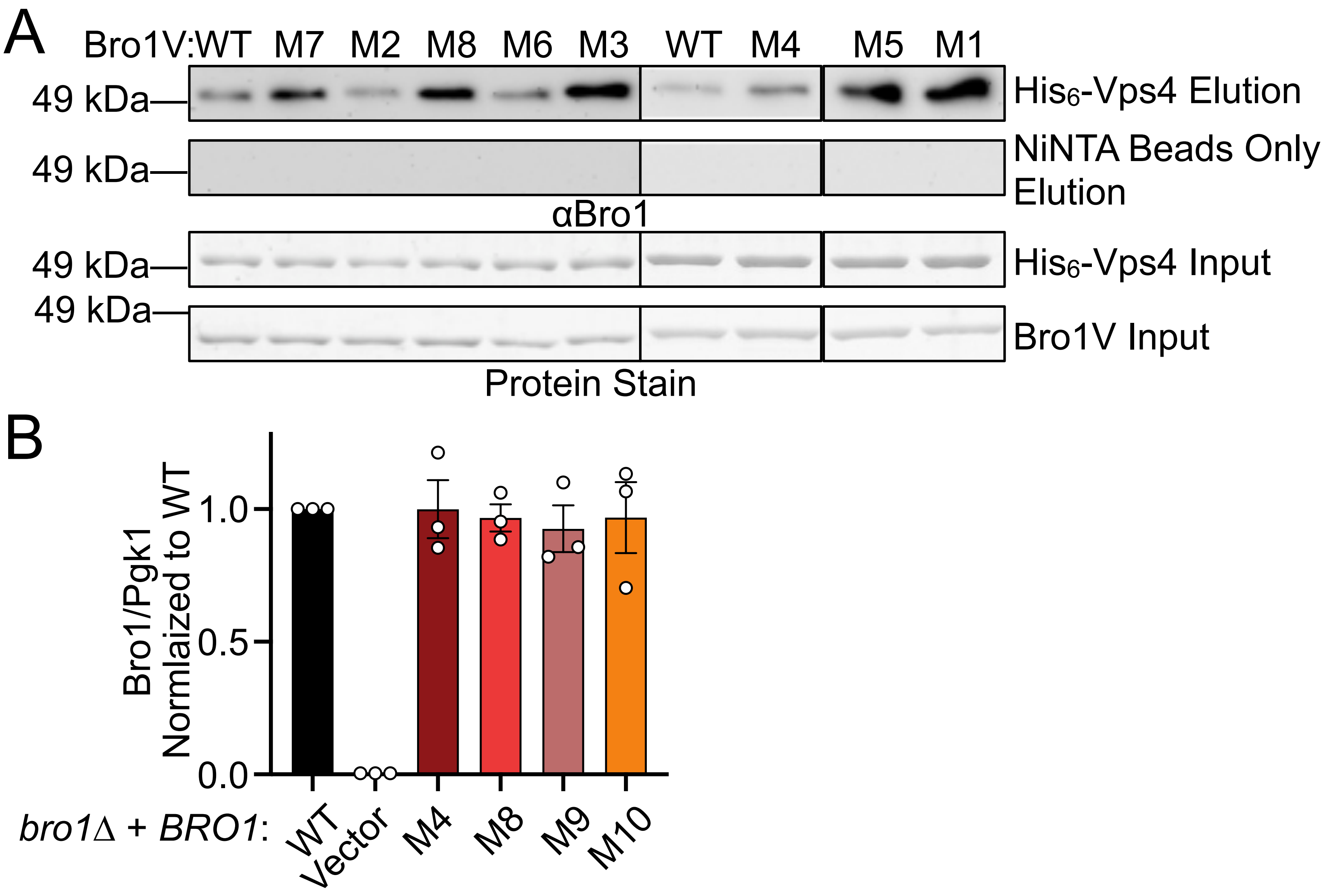

### Supplemental Figure 4

#### Supplemental Figure 4

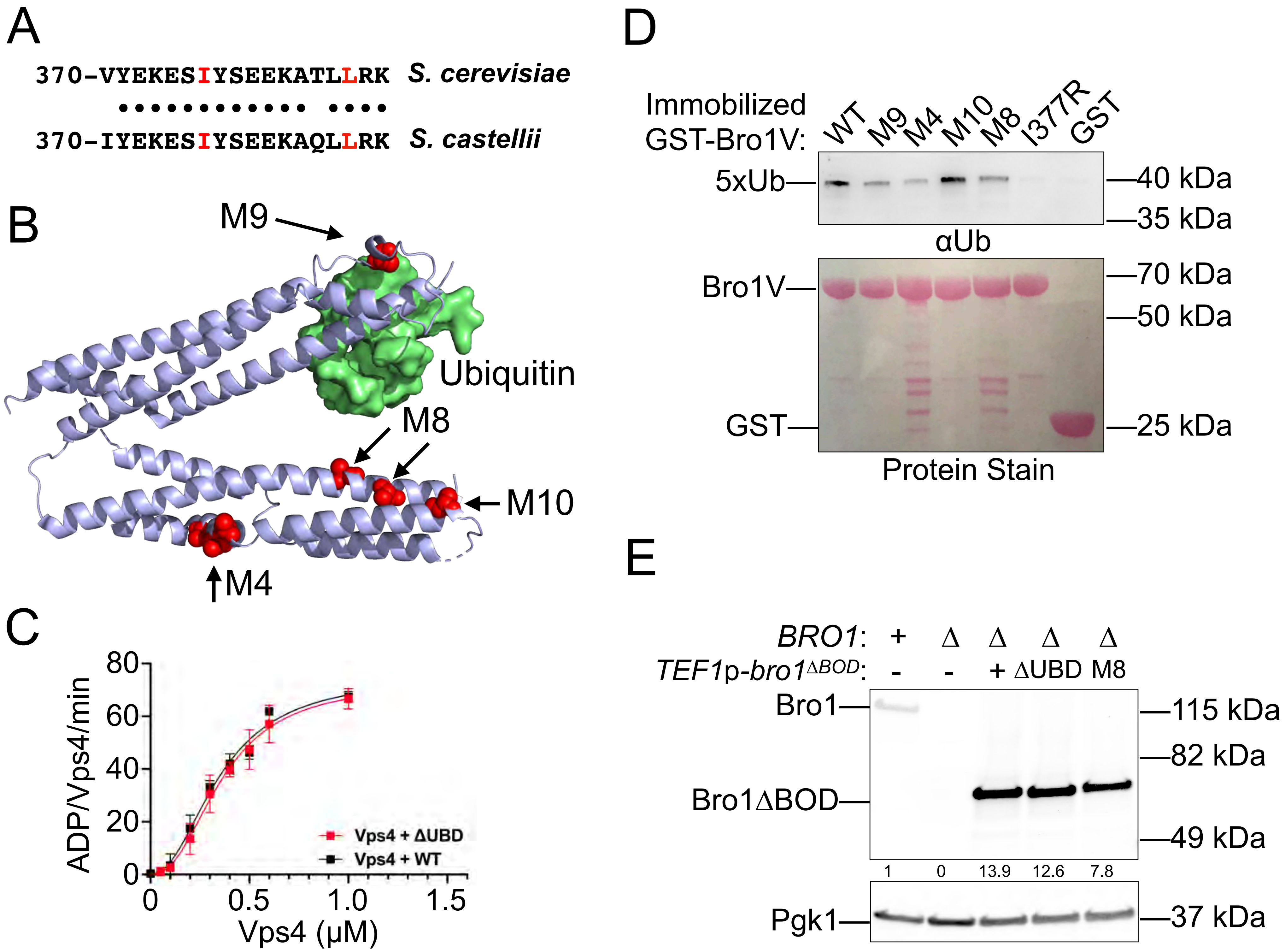
